# Extracellular vesicles improve GABAergic transmission in Huntington’s disease iPSC-derived neurons

**DOI:** 10.1101/2022.12.18.520919

**Authors:** Margarida Beatriz, Ricardo Rodrigues, Rita Vilaça, Conceição Egas, Paulo Pinheiro, George Q. Daley, Thorsten M. Schlaeger, A. Cristina Rego, Carla Lopes

**Affiliations:** CNC-Center for Neuroscience and Cell Biology, CIBB - Centre for Innovative Biomedicine and Biotechnology, University of Coimbra, Coimbra, Portugal; IIIUC-Institute for Interdisciplinary Research, University of Coimbra, Coimbra, Portugal; Biocant- Transfer Technology Association, BiocantPark, Cantanhede, Portugal; Division of Pediatric Hematology/Oncology, Children’s Hospital Boston, Boston, MA USA; Harvard Stem Cell Institute, Boston, MA USA; FMUC-Faculty of Medicine, University of Coimbra, Coimbra, Portugal

**Keywords:** Extracellular vesicles, Huntington’s disease, synaptogenesis, miRNAs

## Abstract

Extracellular vesicles (EVs) carry bioactive molecules associated with various biological processes, including miRNAs. In both Huntington’s disease (HD) models and human samples, altered expression of miRNAs involved in synapse regulation were reported. Recently, the use of EV cargo to reverse phenotypic alterations in disease models with synaptopathy as the end-result of the pathophysiological cascade has become an interesting possibility. Here, we assessed the contribution of EVs to GABAergic synaptic alterations using a human HD model and studied the miRNA content of isolated EVs. After differentiating HD human induced-pluripotent stem cells into electrophysiologically active striatal-like GABAergic neurons, we found that HD-derived neurons displayed reduced density of inhibitory synapse markers and of GABA receptor-mediated ionotropic signaling. Treatment with EVs secreted by control (CTR) fibroblasts reversed the deficits in GABAergic synaptic transmission and increased the density of inhibitory synapses on HD-neuron cultures, while EVs from HD-derived fibroblasts had the opposite effects on CTR-neurons. Moreover, analysis of miRNAs from purified EVs identified a set of differentially expressed miRNAs between manifest HD, premanifest and CTR lines with predicted synaptic targets. The EVs-mediated reversal of the abnormal GABAergic phenotype in HD-derived neurons reinforces the potential role of EVs-miRNAs on synapse regulation.

## Introduction

Huntington’s disease (HD) is an autosomal dominant disorder associated with the expansion of a CAG triplet in exon 1 of the *huntingtin* gene (MacDonald *et al*., 1993). The translated mutant huntingtin (mHTT) has an expanded polyglutamine stretch that causes conformational changes, abnormal protein interactions and protein aggregation (Hoffner *et al*., 2005). The primary disease hallmark is a selective loss of striatal medium spiny neurons (MSN), with symptoms including neuropsychiatric signs and cognitive deficit that can precede motor symptom onset by over 15 years (Beglinger *et al*., 2010; Lemiere *et al*., 2004; Thompson *et al*., 2012). Indeed, neuroimaging studies show that striatal connectivity is altered before clinical diagnosis (Paulsen *et al*., 2014; Tabrizi *et al*., 2012).

Gamma-amino butyric acid (GABA)-ergic MSN, the most common striatal neurons, receive glutamatergic projections from the cortex and their loss typically coexists with loss of cortical pyramidal neurons, primarily in motor and premotor areas (Mehrabi *et al*., 2016; Vonsattel *et al*., 2008). Dysregulation of GABAergic signaling significantly contributes to HD pathogenesis, especially GABA_A_ receptor-mediated signaling, which is altered in the brains of several HD mouse models and in human patients (Hsu *et al*., 2018). MHTT interacts with different organelles and proteins, particularly at synapses, affecting their normal function and ultimately resulting in degeneration (Dargaei *et al*., 2018; Liot *et al*., 2013; Ma *et al*., 2017; Twelvetrees *et al*., 2010; Yuen *et al*., 2012). Both wild type (WT) and mutant forms of huntingtin (HTT) were shown to interact with KCC2, a Cl^-^ cotransporter that promotes GABAergic inhibitory signaling, with a decrease in expression leading to an over-excitation in the hippocampus of an HD mouse model (Dargaei *et al*., 2018). Additionally, mHTT disrupts the translocation of GABA_A_ receptors to the synapse by interfering with the affinity of HTT-associated protein 1 for KIF5, a kinesin motor protein (Twelvetrees *et al*., 2010; Yuen *et al*., 2012). The interaction between mHTT and the SorCS2 protein, that regulates the trafficking of the subunit 2A of NMDA receptors in MSN, leads to motor deficits in a HD mouse model (Ma *et al*., 2017). Furthermore, the dendritic transport of TrkB receptors, that mediate brain-derived neurotrophic factor (BDNF) signaling, is compromised by mHTT, causing diminished activity of the ERK pathway in HD striatal neurons (Liot *et al*., 2013). Moreover, a decrease in GABA content and GABAergic function were measured in the *substantia nigra* of postmortem brain samples of HD patients and in the striatum of HD carriers through PET analysis (Pavese *et al*., 2006; Spokes *et al*., 1980), as well as decreased mRNA and protein levels of GABA_A_R α1 and β2 subunits in the external globus pallidus of HD mouse models (Du *et al*., 2016; Yuen *et al*., 2012). Finally, gephyrin, a scaffold protein present in inhibitory post-synapses, was also decreased in a HD mouse model, along with diminished frequency of miniature inhibitory postsynaptic currents (mIPSC) and GABAergic synapse density (Du *et al*., 2016). Therefore, mHTT can likely act as a disruptor of GABAergic synaptic activity and thus restoring normal GABAergic function could constitute a therapeutic opportunity in HD.

Extracellular vesicles (EVs) are nanosized structures that act as intercellular messengers containing different bioactive cargo, such as lipids, proteins, DNA and miRNAs (Beatriz *et al*., 2022). Recently, the role of miRNAs associated with EVs has been explored in several neurodegenerative diseases either as biomarkers or as modulators of the pathological processes (Beatriz et al.2021). MiRNAs are small, non-coding RNAs that have a modulatory effect on mRNA translation that help maintaining the balance of numerous neuronal mechanisms, including the formation, maturation and function of synapses (Hu and Li, 2017). Importantly, the expression of miRNAs that modulate synaptic function and maturation was altered in the cortex and striatum of HD mouse models (Fukuoka *et al*., 2018; Jin *et al*., 2012). Additionally, the levels of the predicted gene targets were also altered in these animals. One of the downregulated miRNAs was miR-132, involved in the regulation of methyl-CpG binding protein 2 (MeCP2) gene expression that was suggested to abnormally interact with HTT and cause transcriptional dysregulation (Fukuoka *et al*., 2018). *MeCP2* is an important regulator of synaptic plasticity and homeostasis and loss- or gain-of-function mutations trigger opposite changes in synaptic transmission (Na *et al*., 2013). MiR-200a and miR-200c are also altered (Jin *et al*., 2012) and their targets include *neurexin 1* which is important to maintain synaptic function in the striatal circuitry (Davatolhagh and Fuccillo, 2021). In HD postmortem brains, cerebrospinal fluid and plasma, various miRNAs are found altered, including some that are critical for neuronal function (Díez-Planelles *et al*., 2016; Martí *et al*., 2010; Reed *et al*., 2018). Because dysregulated miRNAs may play important roles in HD pathogenesis, a miRNA-based therapeutic strategy could be effective.

An approach with observed effects on counterbalancing pathological alterations is the use of EVs. While the treatment with EVs in HD models has mainly resulted in the observation of reduced mHTT aggregates and increased neuronal survival (Hong *et al*., 2017; Lee *et al*., 2021; Lee *et al*., 2016), studies regarding other pathological conditions reported effects of EV administration on synaptic transmission (Deng *et al*., 2017; Li *et al*., 2020; Long *et al*., 2017; Sharma *et al*., 2019).

In this work, after treating human-derived HD striatal-like neurons with EVs secreted by CTR fibroblasts, we observed improvements on neuronal GABAergic signaling. Contrarily, EVs secreted by HD-derived fibroblasts generated the opposite outcome on CTR-neurons, suggesting a role for EVs on propagation of cellular toxic effects and influencing neuronal function. Moreover, we observed divergent miRNAs profiles in HD- and CTR-EVs, evidencing that EV-miRNAs may play a role as regulators of synaptic-related proteins and be involved in HD pathology.

## Materials and Methods

### Cell culture and reagents

Human dermal fibroblasts were obtained from skin punches of mutant *HTT* gene carriers (Huntington’s disease (HD) carriers) HD carriers and controls (CTRs) as described previously (Lopes *et al*., 2022). A total of nine lines of fibroblasts were used: three CTRs (fCTR1, fCTR2, fCTR3), three premanifest HD (fpHD1, fpHD2, fpHD3) and three manifest HD lines (fHD1, fHD2, fHD3). Fibroblasts were grown in DMEM medium (#D5648, Gibco) supplemented with 9 mM sodium bicarbonate, 10% fetal bovine serum (FBS; Gibco) and 1% penicillin/streptomycin (Gibco). Induced pluripotent stem cells (iPSC) were reprogrammed from human dermal fibroblasts (from two HD carriers derived lines with 43 (fHD3) and 46 (fHD1) CAG repeats and two CTRs (fCTR1, fCTR3; Figure 1D) (Beatriz *et al*., 2022). iPSCs were maintained in Geltrex®-coated plates with StemFlex medium (Gibco). Differentiation into a striatal-specific neuronal phenotype was performed as previously described (Beatriz *et al*., 2022). Briefly, neuronal induction media was supplemented with 5 μM Dorsomorphin (Sigma) and 10 μM SB431542 (Peprotech) until day five and later 1 μM XAV 939 (Peprotech) and 200 ng/ml Sonic Hedgehog (SHH C-25II; R&D) were added until day 12 (Chambers *et al*., 2009; Delli Carri *et al*., 2013; Nicoleau *et al*., 2013). Neuronal progenitors were then dissociated with Accutase (GRiSP) and plated on Geltrex®-coated coverslips in 24-well plates for further neuronal differentiation using N2/B27 medium supplemented with 200 ng/ml SHH, 1μM XAV939 and 30 ng/ml brain-derived neurotrophic factor (BDNF; Peprotech) until day 26. From this point, the medium was changed every three days and only supplemented with 50 ng/ml BDNF to obtain striatal-like neurons at day 80. Cells were maintained in a humidified incubator at 37 °C with 5% CO_2_.

**Figure 1.**
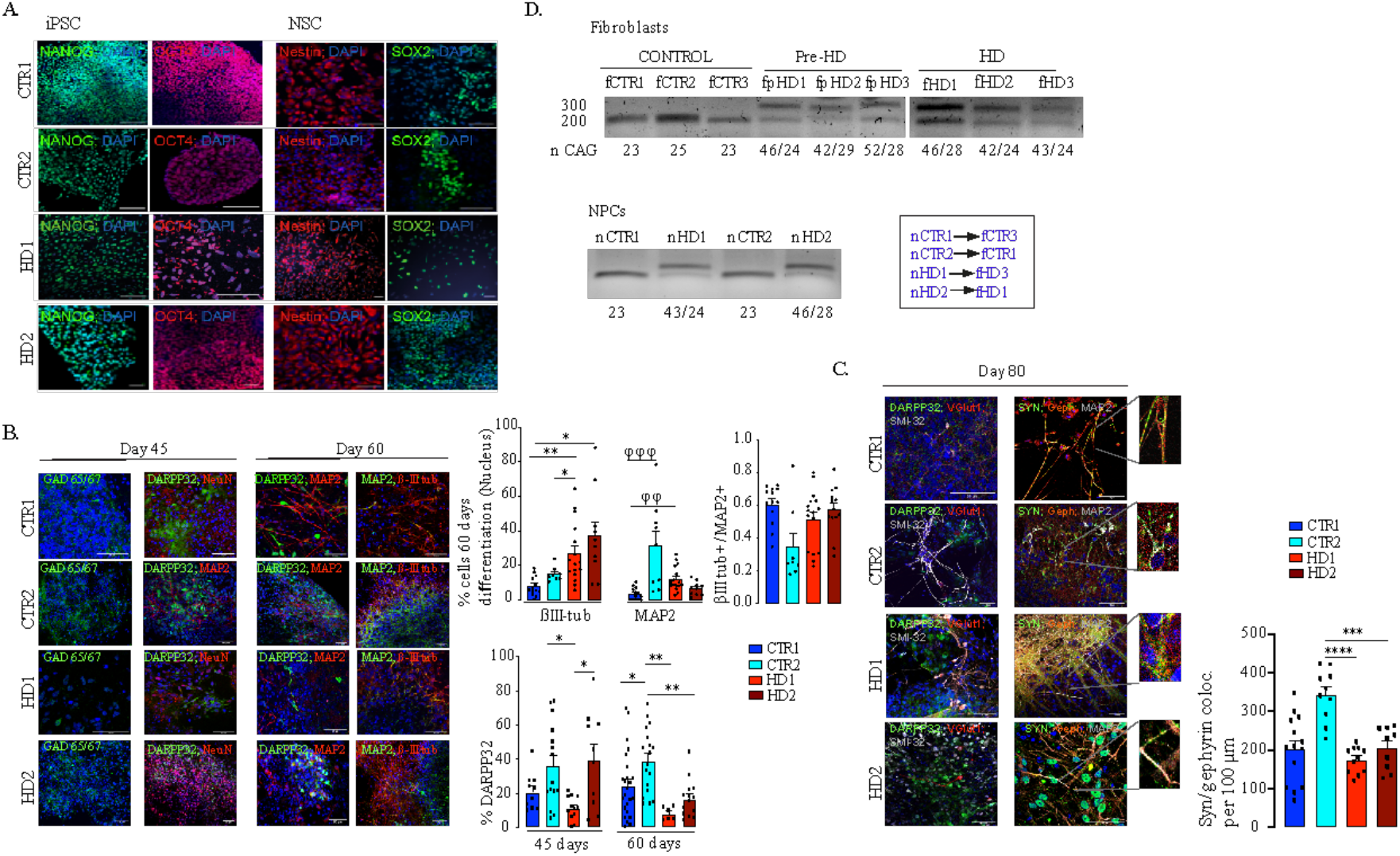
Phenotypic characterization of striatal progenitors and neurons generated from human CTR and HD iPSC. (A) IPSC and differentiated NSC exhibit pluripotency and neural progenitors’ markers (Nanog and OCT4; Nestin and SOX2) (scale bar, 20 μm (B) Neural progenitors express GABAergic synaptic marker (GAD 65/67), neuronal (βIII-tubulin, MAP2 and NeuN) and striatal (DARPP32) specific markers (n>13). βIII-tubulin and MAP2 levels at 60 days and DARPP32^+^ cells during neuronal differentiation at 45 and 60 days (n>10). (C) Differentiated neurons express markers for GABAergic neurons (DARPP32, VGlut1 and SMI-32^+^) and colocalization of synaptic markers for inhibitory (synaptophysin (SYN) with gephyrin (Geph)) showing synapse formation (n>9). (D) CAG repeat sizes for the wild-type (lower band) and mutant alleles (upper band) in fibroblasts and NSC. Correspondence between patients’ fibroblasts and differentiated NSC. Bar plots represent mean±S.E.M. One-way ANOVA followed by Tukey’s multiple comparisons test: * p<0.05, ** p<0.01, *** p< 0.001, **** p< 0.0001; or Kruskalis-wallis followed by Dunn’s multiple comparisons test: φφφ p< 0.001, φφ p< 0.01.

### Immunocytochemistry and image processing

Cells were washed with pre-warmed phosphate-buffered saline (PBS) at 37°C, permeabilized using PHEM buffer (containing: 60 mM PIPES, 25 mM HEPES, 10 mM EGTA and 2 mM MgCl_2_, at pH 6.9) with 0.1% Triton^®^ X-100 and fixed with PHEM/4% paraformaldehyde for 20 minutes. Blocking was performed for 40 minutes using PHEM with 0.1% Triton and 3% bovine serum albumin (BSA). Cells were incubated overnight at 4°C with the following primary antibodies diluted in PHEM/0.1% Triton/3% BSA: anti-NANOG (D73G4; 1:200, #mAb 4903, Cell Signaling), anti-OCT4 (1:200, #2750, Cell Signaling), anti-Nestin (1:200; #MAB1259 R&D), anti-SOX2 (D9B8N; 1:200, #mAb 23064, Cell Signaling), anti-GAD 65/67 (1:100, #ab1511, Merck), anti-DARPP32 (EP720Y; 1:200, #ab40801, Abcam), anti-NeuN (1:500, #MAB377, Chemicon), anti-MAP2 (1:500, #ab92434, Abcam), anti-βIII-tubulin (1:200, #NB100-1612, Novus Biologicals), anti-VGlut1 (1:500, #135 307, Synaptic Systems), anti-SMI-32 (1:500, #837904, BioLegend), anti-Synaptophysin (1:500, #ab32127, Abcam), anti-Gephyrin (1:200, #147111, Synaptic Systems), anti-PSD95 (1:200, #51-6900, Thermofisher), anti-GABA1aR1 (1:50, #224 203, Synaptic Systems) and anti-GFAP (1:500, #AB5804, Chemicon). Cells were washed three times with PBS and further incubated with secondary antibodies diluted in PHEM/0.1% Triton/3% BSA (1:200) for one hour at room temperature (RT) and then washed three times (with a first quick wash and the following two washes for 10 minutes each) with PBS. The coverslips were mounted on glass slides using Mowiol (Sigma); Hoechst (1:1000; Invitrogen) was used to stain the nuclei. Images were acquired on a Zeiss LSM 710 Confocal System (Carl Zeiss Microscopy) with x20 and x40 objectives. Colocalization studies of βIII-tubulin with MAP2 were performed on Z-stacks using the JACoP ImageJ plugin (Bolte and Cordelières, 2006) and synapse counting using the SynapCountJ ImageJ plugin (Mata *et al*.).

### CAG PCR and agarose gel

The confirmation of CAG repeat number in each cell line was determined by PCR using primers for Exon 1 CAG region, as described previously (Evers *et al*., 2015). Briefly, 30 ng of genomic DNA was used for PCR with Phusion® High-Fidelity DNA polymerase in Phusion® GC buffer supplemented with 1M betaine. Amplicons were separated in 2 % agarose gels with TBE (Tris/Borate/EDTA) buffer and visualized with a Bio-Rad Gel Doc XR+ System. A small volume of the reaction was used for a more accurate confirmation of CAG number with Agilent BioAnalyzer. Primers: HttCAG Fw: 5′-ATG GCG ACC CTG GAA AAG CTG AT-3′; HttCAG Rev: 5′-GGC TGA GGC AGC AGC GGC TG-3′.

### Electrophysiological Recordings

Whole-cell patch clamp recordings were performed using an AxoPatch 200B amplifier (Molecular Devices). The borosilicate glass micropipettes used had a resistance of 4–6 MΩ. The voltage-clamp recordings of miniature inhibitory postsynaptic currents (mIPSC) mediated by GABAR were performed at -70 mV using an internal solution containing the following (in mM): CsCl 130, NaCl 10, EGTA 5, HEPES 10, CaCl2 0.5, MgATP 4, NaGTP 0.3, QX314 1 (pH 7.4 with CsOH). The bathing solution contained the following (in mM): NaCl 140, KCl 2.5, CaCl2 2, MgCl2 1, HEPES 10 and glucose 15 (pH 7.4) plus 1 μM tetrodotoxin, 20 μM CNQX and 5 μM 5,7-DiCl-kynurenic acid (Tocris, UK). For whole-cell GABAR-mediated currents, GABA (100 μM), daily prepared in extracellular solution, was rapidly applied to cells with a perfusion valve control system VC-77SP/perfusion fast-step SF-77B (Warner Instruments, USA). The voltage-clamp recordings and the current-clamp recordings of voltage signals were performed using an internal solution containing the following (in mM): K-gluconate 130, NaCl 4, MgCl_2_ 2, EGTA 1, HEPES 10, phosphocreatine 5, Na_2_-ATP 2 and Na_2_-GTP 0.3 (pH 7.2 with KOH). The bathing solution contained the following (in mM): NaCl 140, KCl 3, CaCl_2_ 2, MgCl_2_ 1, HEPES 10 and glucose 15 (pH 7.4). All experiments were performed at RT (22–25°C). The currents were filtered at 1–10 kHz, digitized at a sampling rate of 1-20 kHz to a personal computer and analyzed with pClamp software (AXON Instruments, USA).

### EVs isolation and characterization

Isolation of EVs was performed as previously described (Beatriz *et al*., 2022). Briefly, fibroblasts were cultured in in medium depleted of EVs by centrifuging the FBS at 100 000 x*g* for 18 hours at 4°C. Cell culture supernatant was collected from fibroblasts every 48 hours and stored at -80°C for posterior use. For isolation of EVs, the culture media was filtered (0.22 μm) to remove cell debris, centrifuged at 10 000 *g* for 30 minutes, and twice at 100 000 *g* for 70 minutes to pellet EVs (Théry *et al*., 2006). EVs were resuspended in different buffers depending on the final use: PBS for RNA extraction, Nanosight tracking analysis (NTA) and incubations; PBS/2% paraformaldehyde for transmission electron microscopy (TEM) and RIPA for western blotting.

### TEM and immunogold-TEM

The antibodies anti-1C2 which detects expanded polyglutamine (1:150, #MAB1574, Millipore), and anti-HTT (1:150, #MAB2166, Millipore) were used for immunoelectron labeling of isolated EVs. Briefly, EV samples were placed on carbon-Formvar-coated 300 mesh nickel grids, and samples were allowed to absorb for 20 min. Following washing in PBS (2x 3 minutes), grids were floated (sample side down) onto 50 μl drops of 50 mM glycine (4x 3 minutes) to quench free aldehyde groups and then transferred to blocking solution (5% BSA, 0.01% saponin in PBS) for 10 minutes. Then, grids were incubated in blocking buffer (1% BSA, 0.01% saponin in PBS; negative control) or primary antibodies for two hours, followed by washing steps in PBS with 1% BSA (6x 3 minutes). Secondary antibodies (anti-mouse or anti-rabbit IgC immunogold conjugates, 1:200) were then added for one hour, followed by additional washing steps (8x 2 minutes). After, samples were fixed in 1% glutaraldehyde for five minutes and washed in water (8x 2 minutes). A final contrasting step was performed using uranyl oxalate solution, pH 7.0, for five minutes, followed by a mixture of 4% uranyl acetate and 2% methyl cellulose, for 10 minutes on ice. Imaging was conducted using a FEI-Tecnai G2 Spirit Bio Twin TEM at 100 kV.

### Particle size and concentration analysis

EVs were resuspended in 1 ml of water containing low-mineral concentration. NTA analysis was performed in a NanoSight NS300 instrument with an sCMOS camera module (Malvern Panalytical). Analysis settings were optimized and kept constant between samples. For each sample, five videos were recorded and analyzed to obtain the mean size and estimated concentration of particles. Data were processed using the NTA 3.1 analysis software.

### Western blotting

As previously described (Beatriz *et al*., 2022; Lopes et al., 2020), cells were lysed in a lysis buffer (50 mM Tris-HCl pH 7.4, 150 mM NaCl, 1 mM EDTA, 1% TritonX-100) supplemented with protease inhibitor cocktail (Sigma), and protein was quantified using the Bio-Rad Protein Assay (Bio-Rad). For protein denaturation, SDS sample buffer (50 mM Tris-HCl pH 6.8, 2% SDS, 5% glycerol, 600 mM DTT, 0.01% bromophenol blue) was added to protein lysates (30 μg for fibroblasts) and heated for five minutes at 95°C. For EVs, the final isolated pellet (from 150 ml of fibroblast culture media) was lysed in RIPA buffer and then protein was denatured in SDS sample buffer (non-reducing conditions were used to assess CD63 presence). Samples were then separated on 10-12% polyacrylamide gels and transferred onto methanol-activated polyvinylidene difluoride (PVDF) Hybond-P membranes (Millipore). Membranes were blocked in Tris-buffered saline with 0.1% Tween-20 (TBS-T) containing 5% BSA for one hour at RT and incubated overnight at 4°C with anti-flotillin-2 (C42A3) (1:500, #3436, Cell Signaling), anti-calnexin (1:500, #ab10286, Abcam) and anti-CD63 (TS63) (1:500, #10628D, Life Technologies) diluted in TBS-T/1% BSA. Primary antibodies were washed with TBS-T and membranes were incubated with secondary antibodies anti-mouse and anti-rabbit coupled to alkaline phosphatase (1:5000; Sigma) diluted in TBS-T/1% BSA for one hour at RT. Secondary antibodies were washed with TBS-T, immunoreactive bands were developed with ECF reagent (GE Healthcare) and imaged in a Chemidoc Imaging System (Bio-Rad).

### Incubation of striatal-like neurons with EVs

EVs isolated from fibroblasts (fCTR3 and fpHD3) were diluted in PBS, quantified, and filtered (0.22 μm) as described above before being added to culture media. Approximately, 20×10^6^ EVs were added per well in each incubation (Qadir *et al*., 2018). HD striatal-like neurons were incubated with CTR-EVs and CTR striatal-like neurons with HD-EVs every other day for the last 5 days of differentiation (endpoint 80 days) (Jeon *et al*., 2016). Immunocytochemistry and electrophysiological recordings were made on day 80.

### Plasmids and Viral production

pCL6EGwo and pCL6-CD63eGFP were a kind gift from Dr. Helmut Hanenberg and Dr. Bernd Giebel (University of Duisburg-Essen, Germany). High-titer lentiviral stock particles were produced in HEK293T cells by transiently co-transfecting the described plasmids with psPAX2 (#12260, Addgene) and pCMV-VSV-G (#8454, Addgene) using JetPrime reagent (Polyplus-transfection). After 48 hours, the viral particles were collected from the medium, purified and concentrated as described previously (Tiscornia *et al*., 2006). Lentiviral titer was estimated using FACS based on an encoded fluorescent marker.

### Small RNA Sequencing and Analysis

#### Library preparation and sequencing

Small RNA and miRNA RNA quantity and quality in the nine EV samples were assessed by capillary electrophoresis in the 2100 Bioanalyzer and the Agilent Small RNA kit (Agilent Technologies). miRNA libraries were generated with the NEXTFLEX Small RNA Sequencing Kit v3 (©Bioo Scientific Corp., 2019) from 3-20 ng of small RNA for each sample. NEXTFLEX 3’ 4N Adenylated adapter and NEXTFLEX 5’ 4N Adenylated adapter were diluted 1:4 as recommended for low input library preparation. NEXTFLEX tRNA/YRNA Blockers (Bioo Scientific, Perkin Elmer, Waltham, MA) were applied to deplete the tRNA and YRNA fragments. To increase yield, the 3’ adapter ligation was performed at 20ºC, overnight. First strand cDNA synthesis was obtained by reverse transcription for 30 min at 42ºC followed by 10 min at 90ºC. Second Strand cDNA was obtained by PCR amplification for 25 cycles, with a Universal and a barcoded primer (one for each sample), included in the NEXTFLEX Small RNA Sequencing Kit v3. A PAGE-based size selection method was used for obtaining a purified ∼150bp miRNA library. All library preparation procedures were carried out according to manufacturer’s instructions. Sequencing was performed at Genoinseq (Cantanhede, Portugal) on a NextSeq 550 Illumina sequencer with the NextSeq 500/550 MID Output (150 cycles) v2.5 kit (Illumina).

#### Data processing

Sequence data was processed at Genoinseq (Cantanhede, Portugal). Raw single-end reads (R1) were extracted from Illumina Nextseq® System in fastq format and quality-filtered with Trimmomatic version 0.3 (Bolger *et al*., 2014) and Prinseq version 0.20.4 (Schmieder and Edwards, 2011) to clip the NEXTFLEX Small RNA Sequencing Kit adapter and the first and last 4 N bases introduced during library preparation and to remove reads shorter than 18 bp. Ribosomal and transfer RNA reads were removed from the high quality data by alignment against the RFam version 14.5 (Kalvari *et al*., 2021) using Bowtie version 1.0.0 (Langmead *et al*., 2009). The miRDeep2 package version 2.0.1.3 (Friedländer *et al*., 2012) was used to profile known miRNA expression in the 9 samples. The cleaned sequencing data was mapped against the human reference genome assembly GRCh37 (hg19) with Bowtie version 1.0.0, using default parameters and without mismatches. Then the mapped reads were aligned against the miRBase version 22 (Kozomara *et al*., 2019), filtered for human miRNAs only, to identify and quantify all known mature miRNAs and their precursor sequences, also without allowed mismatches. miRDeep2 was also used to predict novel miRNAs from unannotated reads.

The sequencing, filtering and alignment metrics are presented in supplementary Table 1.

#### EVs miRNA target prediction and pathway enrichment analysis

miRNA raw read counts were divided by the total count of their sample for normalization. To identify differentially expressed miRNA we applied an FDR-adjusted p-value (q ≤ 0.05) and fold changes (log2FC ≥ |2.0|). Fold-change was defined as the ratio (log2) between the average expression of the three samples for each two analyzed conditions (CTR, pHD and HD). miRDB-MicroRNA Target Prediction Database (http://mirdb.org/) was used to retrieve the EV miRNA gene targets (Chen and Wang, 2020). Gene Ontology (GO) enrichment analyses for candidate miRNA targets were performed by using the online Web-based tool NetworkAnalyst (https://www.networkanalyst.ca/) (Xia *et al*., 2015; Zhou *et al*., 2019). NetworkAnalyst supports enrichment analysis with gene sets from the Gene Ontology, PANTHER among other databases. SynGo was used to analyze the brain-expressed genes (https://syngoportal.org/) (Koopmans *et al*., 2019). The gene ontology enrichment analysis was performed for the category GABA-ergic synapse (GO:0098982). A normalized average gene expression of SynGO genes (release 20210225-1.1) in GABAergic neurons obtained from a total number of 177614 GABAergic cells and of 1205 SynGO genes in the dataset, kindly provided by Dr Verhage and Dr Roig Adam, was used. Principal Component Analysis (PCA) plots and heatmaps was performed in the web tool ClustVis (https://biit.cs.ut.ee/clustvis/) (Metsalu and Vilo, 2015). Venn diagrams were made at https://bioinformatics.psb.ugent.be/webtools/Venn/.

### Statistical analysis

Statistical computations were performed using GraphPad Prism version 9.0, GraphPad Software, La Jolla, CA, USA, and SPSS version 21.0 (IBM SPSS Statistics for Windows, IBM Corp). Results are represented as the mean ± SEM of the indicated number of independent experiments in figures legends. For cell experiments, at least three independent assays were performed for each experimental condition. Statistical significance was analyzed using parametric test, one-way ANOVA, followed by Bonferroni post hoc test and non-parametric test Kruskal Wallis followed by Dunn’s multiple comparison test. Correlations were done using the Spearman rank correlation coefficient (ρ). p<0.05 or q(p-adjusted) <0.05 was considered significant.

## Results

### Differentiation and maturation of striatal-like neurons

In this study, we used a previously published protocol (Beatriz *et al*., 2022; Delli Carri *et al*., 2013; Lopes et al., 2020) to promote neural induction in pluripotent cell colonies and obtain mature neurons (Figure S1A). Pluripotency (Nanog and OCT4) and neural induction (Nestin) marker were assessed in iPSCs and NSCs, respectively (Figure 1A). During the differentiation protocol, cells were characterized for the presence of markers of GABAergic (GAD 65/67^+^) and MSN (DARPP32^+^) development, and for commitment to a mature neuronal fate by assessing βIII-tubulin^+^ and MAP2^+^ cell populations on days 45 and 60 of differentiation (Figure 1B). The HD cell lines have the capacity to mature at the appropriate time course. Dense clusters of βIII-tubulin^+^ and MAP2^+^ cells are present after 60 days of differentiation, despite an increase in βIII-tubulin^+^ reactivity in HD cells (Figure 1B). Furthermore, when analyzing the expression of DARPP32 to identify the capacity for terminal maturation into MSN, the HD-derived neuronal cultures show a reduction overtime (from 45 to 60 days) compared to CTRs (Figure 1B). By day 80, the majority of differentiated cells expressed the axonal marker SMI-312, but very few neurons were immunoreactive for VGlut1 (Figure 1C). To assess the successful establishment of inhibitory synapses, we measured inhibitory synapse density based on the colocalization of a general presynaptic marker protein, synaptophysin, with the inhibitory postsynaptic protein Gephyrin. We found that synapse density was decreased in HD striatal-like neurons compared to CTR2 (Figure 1C). We also assessed excitatory synapse density through the colocalization of synaptophysin with the excitatory postsynaptic protein PSD-95 and found to be decreased in HD1-neurons compared to CTR2 and in HD2 compared to both CTRs (Figure S1B). In addition, we observed the presence of a GFAP^+^ glial population of cells (Figure S1C). The abnormal CAG expansion in HD-derived cells was confirmed by PCR (Figure 1D). Together, these data demonstrate that HD-derived neurons show a potential delay in maturation into an MSN phenotype and display altered synapse density.

To further validate a delayed maturation from a functional perspective, the electrophysiological properties of differentiated HD and CTR neurons were analyzed through whole-cell current-clamp recordings. At day 60 of the differentiation protocol, neurons were still immature and action potential firing capacity, upon injection of depolarizing current pulses, was low. However, at day 80, both HD- and CTR-derived neurons were capable of generating action potentials in response to depolarizing current injections, exhibiting single and repetitive spiking, indicating similar levels of maturation (Figure 2A). The mean resting membrane potential of HD striatal-like neurons was shifted toward more positive values (HD1: 42.2 ± 2.0 mV and HD2: -33.7 ± 2.0 mV) compared to CTRs (CTR1: -46.9 ± 1.9 mV and CTR2: -48.6 ± 1.6 mV) indicating that HD neurons are more depolarized, although only HD2 presented statistical significance (p<0.05). Nonetheless, the action potential threshold was the same between HD and CTR cells (Figure 2B). We also found that HD striatal-like neurons had a lower rheobase compared to CTR2 neurons, which reflects a greater excitability (Figure 2B). In agreement with this, the HD2 cell line also exhibited a higher input resistance compared to the CTR neurons (Figure S2A). Since voltage-gated sodium (Na^+^) and potassium (K^+^) channels are essential players in neuronal physiology, we evaluated the activation of fast voltage-gated Na^+^ and delayed-rectifier K^+^ currents (Figure 2C). The normalized I/V curves for Na^+^ suggest that the activation threshold is similar between CTRs and HDs despite the latter having a tendency for larger maximum current densities, probably due to differences in the membrane levels of Na^+^ channels. The I/V curves for K^+^ show a slight shift in activation toward a more positive voltage in the HD2 cell line. No significant alterations were observed between HDs and CTRs for maximum K^+^ current density (Figure 2C).

**Figure 2.**
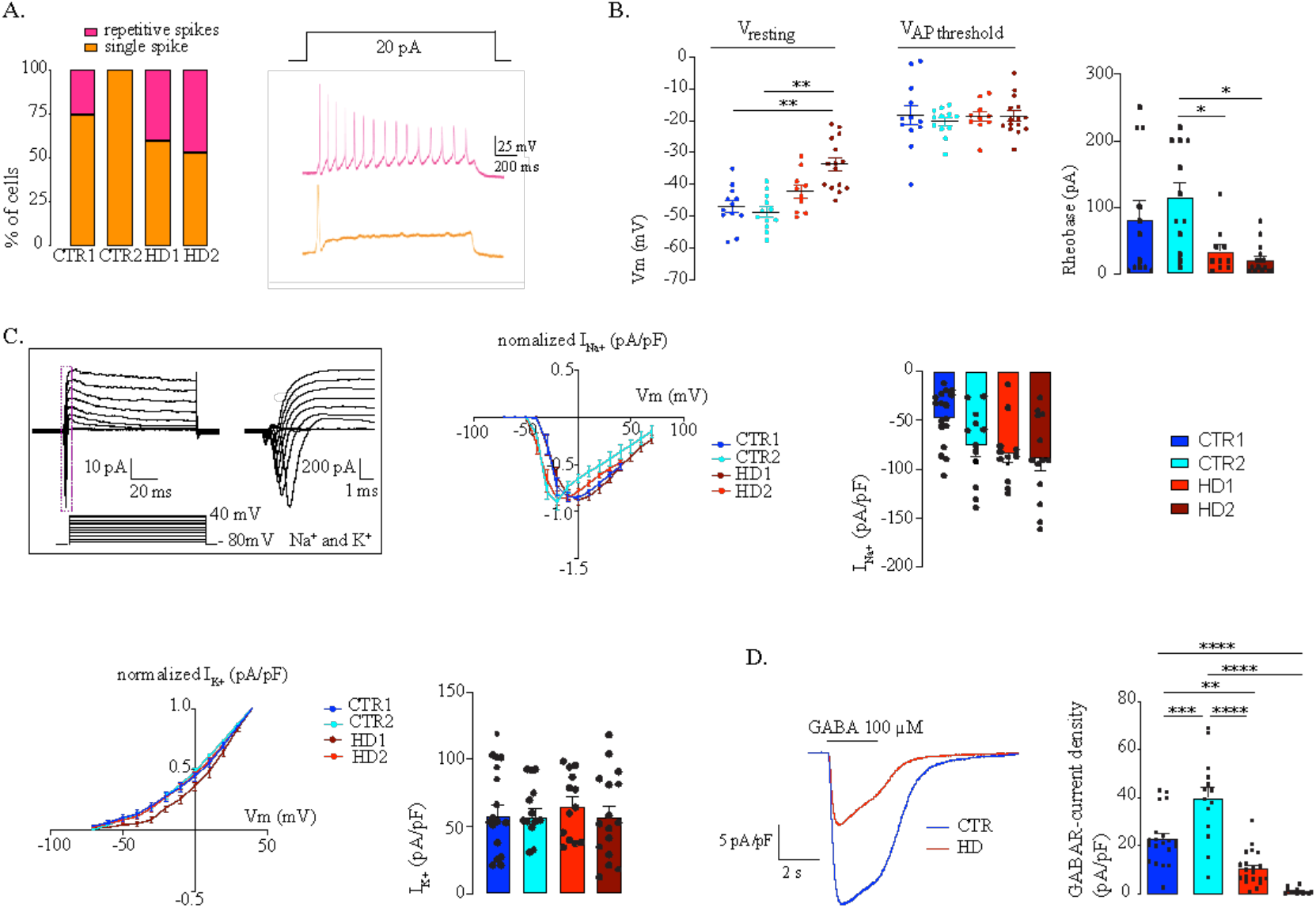
Electrophysiological characterization of mature striatal-like neurons. (A) Differentiated neurons are able to generate single or sustained action potentials in response to depolarizing current injections. B) HD-derived neurons have a less negative resting potential. C) HD-derived neurons have more depolarized resting membrane potential but unaltered action potential threshold and display altered voltage-dependence of Na^+^ and K^+^ conductance. D) HD-derived neurons show reduced GABA_A_ receptor-mediated currents in response to rapid application of GABA (100 μM). Bar plots represent mean±S.E.M. One-way ANOVA followed by Tukey’s multiple comparisons test: * p<0.05, ** p<0.01, *** p< 0.001, **** p< 0.0001.

Since we aimed to obtain an enriched population of GABAergic neurons, we further analyzed if GABA receptors were expressed by using a fast bath application of 100 μM GABA and measured the GABA receptor-mediated currents. The whole cell GABA receptor-mediated current density was significantly smaller in HD-striatal-like neurons compared to CTRs (CTR1: 22.9 ± 2.4 pA/pF; CTR2: 39.8 ± 4.7 pA/pF; HD1: 10.4 ± 1.5 pA/pF and HD2: 1.3 ± 0.4 pA/pF; p<0.05) (Figure 2D), indicative of an overall lower expression of GABA receptors. Collectively, these data show that differentiated neurons presented an MSN phenotype and that HD-derived neurons were more excitable and had a decreased GABA receptor-mediated current density.

### HD-neurons display a lower density of inhibitory synapses that is normalized upon EVs incubation

It has been demonstrated that EVs carry miRNAs that are important for cell signaling pathways including those known to promote synaptogenesis (Upadhya *et al*., 2020). To evaluate the effects of EVs on mature striatal-like neurons, we first isolated small EVs and characterized them according to ISEV guidelines (Théry *et al*., 2018). EVs were isolated from the cell culture media of human fibroblast lines from three CTR donors and six HD carriers (premanifest and manifest) (Lopes *et al*., 2022) using a multi-step centrifugation protocol (Théry *et al*., 2006). EVs presented the expected cap-shaped morphology when observed by TEM (Figure 3A) and a size range of 50-200 nm (Figure 3B) (Théry *et al*., 2018). Small EV protein markers like Flotillin-2 and CD63 were present; the absence of Calnexin as a cellular contaminant validated the purity of the isolated pool of vesicles (Figure 3C). As shown previously (Jeon *et al*., 2016; Kuang *et al*., 2021), we also detected the presence of both HTT and mHTT in EVs from fibroblasts of one CTR and two HD carriers (premanifest and manifest), respectively, by immunogold-TEM (Figure 3D).

**Figure 3.**
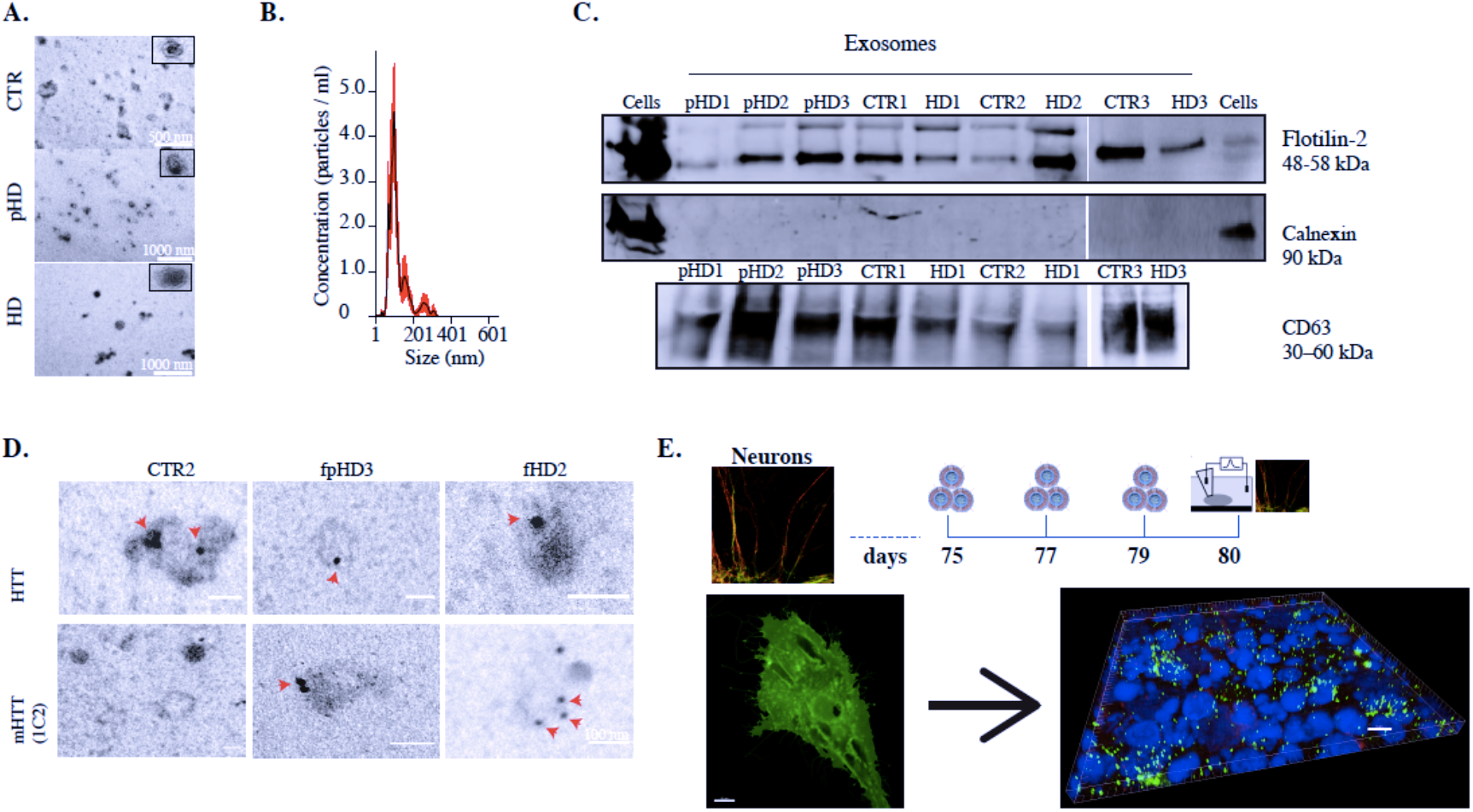
Characterization of fibroblasts derived-EVs. (A) TEM images of fibroblasts-EVs (scale bar, (fCTR2) 500 nm, (fpHD3; fHD2) 1000 nm). (B) Representative analysis of NTA measurements of fibroblasts-EVs concentration (particles/ml) and size (nm). (C) Representative Western blot with conventional EVs markers in fibroblasts-EVs (Flotillin-2 and CD63) and Calnexin as a negative control for cellular contamination. (D) HTT and mHTT (1C2) immunogold labeling in fibroblasts-EVs (scale bar, 100 nm). (E) Timeline for incubation of CTR and HD differentiated neurons with EVs isolated from CD63+-GFP expressing fibroblasts; 3-D reconstruction with IMARIS software of the Z -stacks of confocal images (EVs-green, MAP2-red) Scale, 50 μm.

To assess the effects of EVs in differentiated striatal-like neurons, we applied an incubation protocol of fibroblasts-EVs (using EVs secreted from one CTR and one HD carrier fibroblast line) during the last five days of neuronal differentiation (Figure 3E) (Jeon *et al*., 2016; Qadir *et al*., 2018). Filtered EVs were added to neurons every other day until day 80 of differentiation. To confirm the uptake of EVs by neuronal cultures, we performed a parallel incubation of neuron cultures with EVs obtained from GFP-CD63-expressing fibroblasts. 3D reconstruction with IMARIS software shows EVs inside MAP2^+^ neurons (Figure 3E). After incubation of HD-derived neurons with CTR-EVs for five days, a normalization in the GABAR-mediated current density was observed, whilst incubation of CTR-neurons with HD-EVs resulted in the opposite effect (Figure 4A).

**Figure 4.**
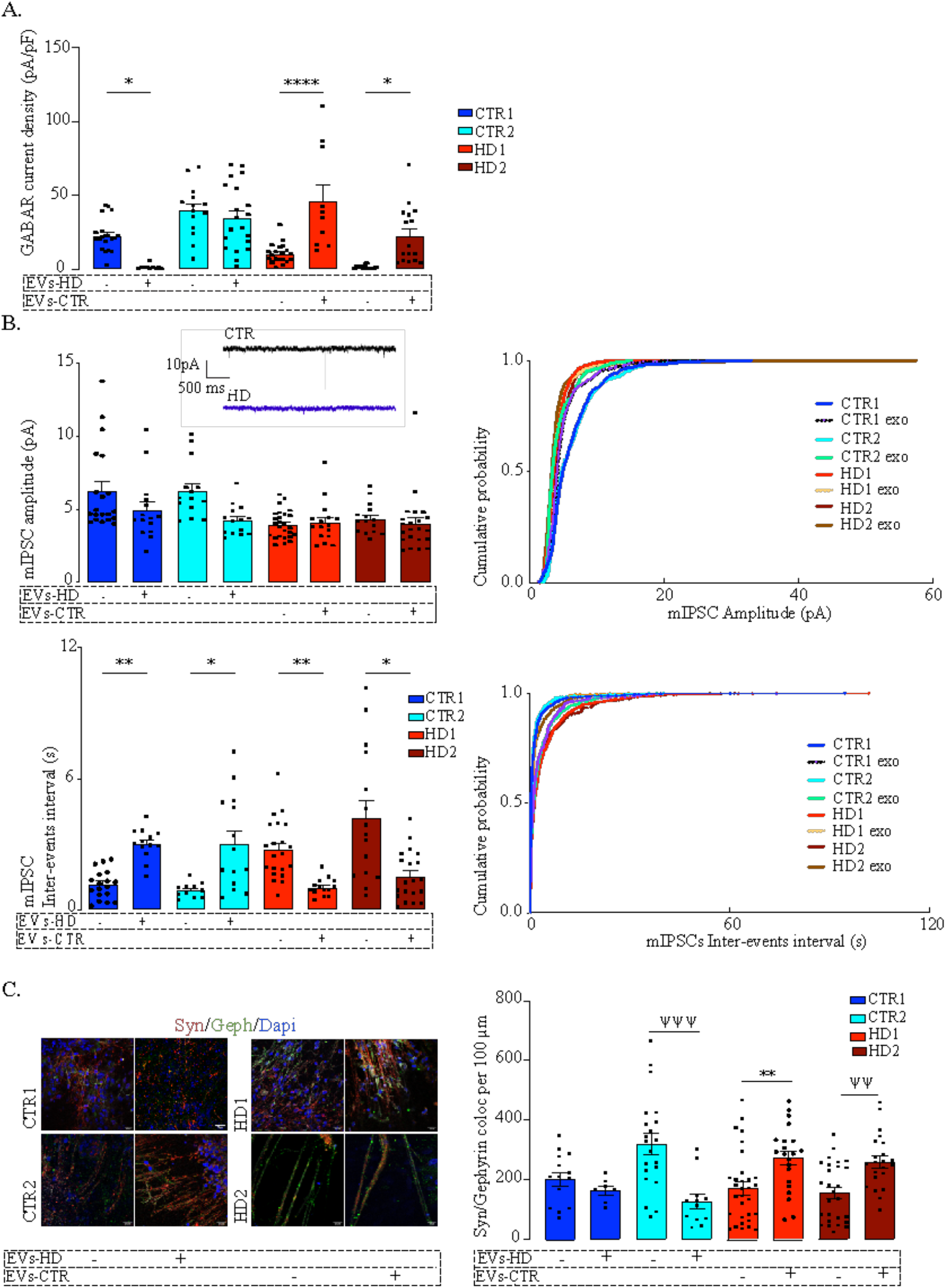
Fibroblasts derived-EVs ameliorate abnormal GABAergic function in human HD iPSC-Derived Neurons. (A) HD-derived neurons reduced GABA_A_ receptor-mediated currents are rescued by incubation with EVs from CTR fibroblasts. (B) HD-derived neurons display reduced mIPSC amplitude and decreased miniature inhibitory postsynaptic current (mIPSC) frequency (increased inter-event interval) and incubation with CTR-EVs recovered only the frequency of events in HD-derived neurons while the opposite effects occur in CTR-derived neurons incubated with HD-EVs. (C) Increased colocalization of synaptic markers for inhibitory (synaptophysin-red with gephyrin-green) in HD-derived neurons upon incubation with EVs from CTR fibroblasts indicating enhanced synapse formation; the opposite effect was observed in CTR-derived neurons exposed to HD fibroblasts EVs. Scale, 20 μm. Bar plots represent mean±S.E.M. One-way ANOVA followed by Tukey’s multiple comparisons test: * p<0.05, ** p<0.01, **** p< 0.0001; or Mann-Whitney: ψψ p<0.01, ψψψ p< 0.001.

Next, we assessed if, in addition to having surface GABA receptors, the differentiated neurons have synaptic GABA receptor-mediated mIPSC and whether these are influenced by treatment with EVs. HD-derived neuron lines had lower mIPSC frequency (higher inter-event intervals) that increased significantly after incubation with CTR-EVs, suggesting an increase in the number of active synaptic contacts. No differences were found in mIPSC amplitude between non-treated neurons and upon EVs treatment, suggesting that the number of GABA receptors at individual synapses was unchanged (Figure 4B). To confirm that EVs were causing an increase in the number of synaptic contacts, we quantified the colocalization of Synaptophysin with Gephyrin as a measure of GABAergic synapse density. In the HD-derived neurons treated with CTR-EVs a significant increase in inhibitory synapse density was observed, while in CTR-derived neurons treated with HD-EVs the opposite effect occurred (Figure 4C and S2). Therefore, while EVs secreted by CTR fibroblasts promote synapse formation, EVs secreted by HD-derived fibroblasts impair this process.

### EVs are enriched in miRNAs targeting synaptic genes

To explore the hypothesis that miRNAs carried by EVs could be involved in the synaptogenic effect, we analyzed the miRNAs profile of EVs released from the three CTRs and six HD (three premanifest-pHD1, pHD2, pHD3- and three manifest-HD1, HD2, HD3) fibroblast lines used for the previous incubations. In total, 392 miRNAs were detected in EVs, and two distinct miRNA clusters could be observed after hierarchical grouping of the top 100 miRNAs found (CTRs, pHDs and HDs) although samples did not cluster distinctly according to their origin (Figure 5A). The principal component analysis (PCA) showed a clear distinction between EVs isolated from premanifest patients and CTRs, except for the CTR2 cell line (Figure 5B). When looking at miRNA composition, 227 miRNAs were detected across all group samples (Figure 5C, D). Once analyzing only for premanifest and manifest lines, we observed the smallest number of EV-associated miRNAs in common (8) when compared to premanifest vs CTR (20) or manifest vs CTR (32) (Figure 5E). Interestingly, some of the miRNAs were exclusively detected in a specific cell derived EV population, confirming the selective incorporation of miRNAs into EVs (Figure 5C).

**Figure 5.**
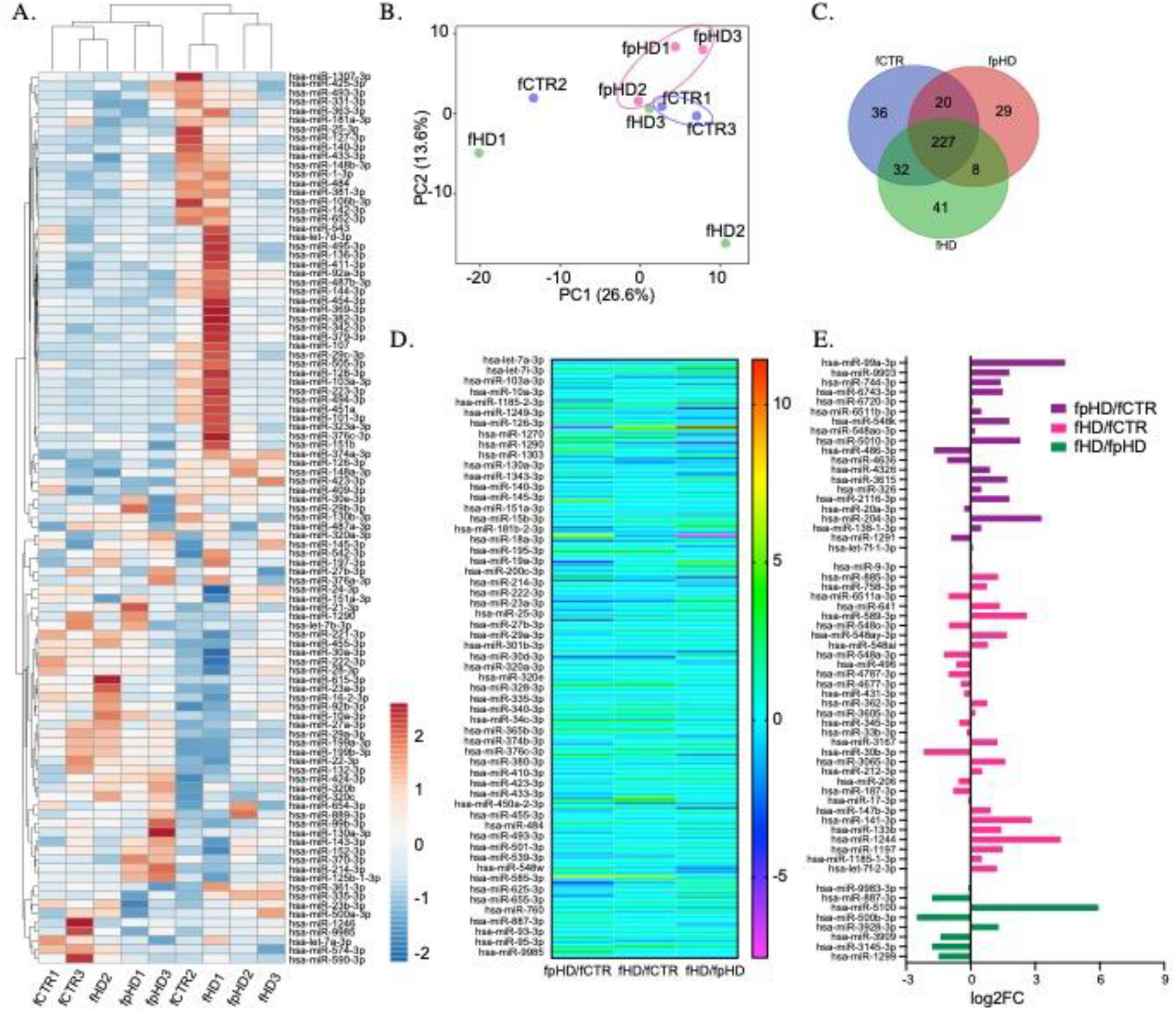
Characterization of miRNAs in extracellular vesicles released from HD and CTR fibroblasts. (A) Heatmap showing the most abundant 100 miRNAs in EVs isolated from the nine fCTR, fpHD and fHD cell lines. High expression based on raw read counts normalized using linear transformations is shown in red while low expression is shown in blue. Rows and columns are clustered using correlation distance and average linkage. 100 rows, 9 columns. (B) PCA showing EVs miRNA profiles for each group (n=9 data points); X and Y axis show principal component 1 and principal component 2 that explain 26.6% and 13.6% of the total variance, respectively. (C) Venn diagram showing the number of miRNAs with specific expression for each group (fCTR, fpHD and fHD lines). (D) Heatmap generated from EVs-miRNAs comparing fold changes in the levels of the 206 miRNAs common to all groups. Heatmap colors represent gene expression fold change levels based on the provided color key scale; red for upregulated gene expression levels, pink for downregulated gene expression levels. (E) Fold change for differentially expressed genes common to the two groups according to Venn diagram.

Since we could not identify differentially expressed miRNAs between samples (q-value <0.05), probably due to the reduced sample size, we further analyzed miRNA by their fold change. From the upregulated miRNAs with logFC > 2 we found 28 for pHD vs CTR; 23 for HD vs CTR and 18 for HD vs pHD. The downregulated miRNAs with logFC > -2 were less numerous: 12 for pHD vs CTR; 2 for HD vs CTR and 13 for HD vs pHD (table 1, suppl. table 1). Among the most enriched miRNAs in EVs, hsa-miR-576-3p was upregulated in both HD and pHD vs CTRs, hsa-miR-1260a was found upregulated in HD and downregulated in pHD vs CTRs and hsa-miR-1260b was more abundant in HD-EVs than in the other groups (table 1, suppl. table 1).

**Table 1.** MicroRNAs up- and down-regulated in fibroblasts-derived EVs from patients with HD

| upregulated |  |  | downregulated |  |
| --- | --- | --- | --- | --- |
|  | miRNA | FDR | miRNA | FDR |
| pHD/CTR | hsa-miR-576-3p* | 6,34 | hsa-miR-1260a | -5,95 |
|  | hsa-miR-184 | 6,25 | hsa-miR-19a-3p | -4,61 |
| | hsa-miR-99a-3p | 4,40 | hsa-miR-30d-3p \$ | -3,90 |
|  | hsa-miR-34c-3p* | 4,14 | hsa-miR-590-3p | -2,95 |
|  | hsa-miR-210-3p | 3,82 | hsa-miR-26b-3p | -2,79 |
|  | hsa-miR-1277-3p* | 3,70 | hsa-miR-2355-3p | -2,56 |
|  | hsa-miR-377-3p | 3,67 | hsa-miR-18a-3p | -2,49 |
|  | hsa-miR-195-3p* | 3,66 | hsa-miR-598-3p | -2,44 |
|  | hsa-miR-1228-3p | 3,54 | hsa-miR-2110 | -2,24 |
|  | hsa-miR-129-2-3p | 3,51 | hsa-miR-6529-3p | -2,17 |
|  | hsa-miR-4454* | 3,46 | hsa-miR-1180-3p | -2,15 |
|  | hsa-miR-548l* | 3,40 | hsa-miR-181a-3p | -2,12 |
|  | hsa-miR-204-3p | 3,31 |  |  |
|  | hsa-miR-940 | 2,83 |  |  |
|  | hsa-miR-323b-3p* | 2,59 |  |  |
|  | hsa-miR-34a-3p* | 2,41 |  |  |
|  | hsa-miR-5010-3p | 2,29 |  |  |
|  | hsa-miR-324-3p | 2,28 |  |  |
|  | hsa-miR-3158-3p | 2,27 |  |  |
|  | hsa-miR-10401-3p | 2,25 |  |  |
|  | hsa-miR-1843 | 2,21 |  |  |
|  | hsa-miR-1185-2-3p | 2,21 |  |  |
|  | hsa-miR-365a-3p* | 2,18 |  |  |
|  | hsa-miR-365b-3p* | 2,18 |  |  |
|  | hsa-let-7e-3p | 2,13 |  |  |
|  | hsa-miR-548w | 2,11 |  |  |
|  | hsa-miR-339-3p | 2,05 |  |  |
|  | hsa-miR-133a-3p | 2,04 |  |  |
| HD/CTR | hsa-miR-576-3p | 5,83 | hsa-miR-30b-3p | -2,18 |
| | hsa-miR-1260a # | 5,48 | hsa-miR-30d-3p \$ | -3,34 |
|  | hsa-miR-4429 # | 4,96 |  |  |
|  | hsa-miR-1260b # | 4,62 |  |  |
|  | hsa-miR-1244 | 4,19 |  |  |
|  | hsa-miR-195-3p | 3,96 |  |  |
|  | hsa-miR-1277-3p | 3,91 |  |  |
|  | hsa-miR-4454 | 3,57 |  |  |
|  | hsa-miR-34c-3p | 3,39 |  |  |
|  | hsa-miR-323b-3p | 3,17 |  |  |
|  | hsa-miR-141-3p | 2,84 |  |  |
|  | hsa-miR-98-3p | 2,80 |  |  |
|  | hsa-miR-376b-3p # | 2,75 |  |  |
|  | hsa-miR-589-3p | 2,61 |  |  |
|  | hsa-miR-450a-2-3p # | 2,54 |  |  |
|  | hsa-miR-548l | 2,45 |  |  |
|  | hsa-miR-412-3p | 2,37 |  |  |
|  | hsa-let-7i-3p | 2,36 |  |  |
|  | hsa-miR-122-3p # | 2,29 |  |  |
|  | hsa-miR-34a-3p | 2,28 |  |  |
|  | hsa-miR-365a-3p | 2,14 |  |  |
|  | hsa-miR-365b-3p | 2,14 |  |  |
|  | hsa-miR-296-3p | 2,06 |  |  |
| HD/pHD | hsa-miR-1260a | 11,43 | hsa-miR-184 | -7,54 |
|  | hsa-miR-5100 | 8,80 | hsa-miR-129-2-3p | -4,98 |
|  | hsa-miR-4429 | 6,07 | hsa-miR-210-3p | -4,84 |
|  | hsa-miR-19a-3p | 5,05 | hsa-miR-1228-3p | -3,59 |
|  | hsa-miR-1260b | 3,58 | hsa-miR-940 | -2,57 |
|  | hsa-miR-6529-3p | 3,26 | hsa-miR-500b-3p | -2,53 |
|  | hsa-miR-23c | 3,13 | hsa-miR-133a-3p | -2,38 |
|  | hsa-miR-122-3p | 3,02 | hsa-miR-30c-1-3p | -2,37 |
|  | hsa-miR-19b-3p | 2,88 | hsa-miR-1270 | -2,33 |
|  | hsa-miR-483-3p | 2,47 | hsa-miR-4510 | -2,28 |
|  | hsa-miR-1180-3p | 2,38 | hsa-miR-1304-3p | -2,15 |
|  | hsa-miR-671-3p | 2,33 | hsa-miR-377-3p | -2,14 |
|  | hsa-miR-1303 | 2,19 | hsa-miR-708-3p | -2,03 |
|  | hsa-miR-26b-3p | 2,16 |  |  |
|  | hsa-miR-450a-2-3p | 2,16 |  |  |
|  | hsa-miR-376b-3p | 2,12 |  |  |
|  | hsa-miR-585-3p | 2,07 |  |  |
|  | hsa-miR-590-3p | 2,02 |  |  |

Data are shown as fold-change relative to control group
\* Overexpressed miR in commom between pHD/CTR and HD/CTR

To identify the putative genes targeted by these miRNAs, we used miRDB, an online database, to predict the target genes of miRNAs that were up- or downregulated at least two-fold in our EV populations (suppl. table 2). The prediction results were sorted in descending order, as ranked by the target score, and only candidate transcripts with scores ≥ 60 are presented, to increase the prediction confidence (suppl. table 3).

We next performed a target enrichment analysis to identify cellular pathways or functional categories of the putative target genes. By analyzing the biological process, we found that the target genes of the upregulated miRNAs in EVs were mostly related to “Regulation of transcription by RNA polymerase II” or “Chromatin organization”. For the cellular component, these proteins were enriched in “Nucleus”, “Cytoplasm” or “Intracellular” (suppl. table 4). The target genes of the downregulated miRNAs were mostly associated to the cellular processes “Regulation of transcription by RNA polymerase II”, “Cell cycle”, “Apoptotic process”, “Protein phosphorylation” or “amino acid biosynthetic process”. For the cellular component, these genes were enriched in “nucleus”, “cytoplasm”, “Plasma membrane” or “Intracellular” (suppl. table 3). Interestingly, we observed an enrichment for targets in neuronal and synaptic components in both up- and downregulated miRNAs (suppl. table 4). Moreover, the target genes of the highly enriched miR- 1260 family in HD-EVs included the cellular components “Neuronal cell body”, “Dendrite”, “Axon” and “Presynaptic membrane”, showing they can potentially target neuronal activity. Also, the target genes of the upregulated hsa-miR-576-3p in the EVs from HD and pHD show an enrichment profile for the cellular components “Neuron projection” and “Axon” (fig S3; suppl. table 4).

To confirm the synaptic functions of the target genes we used the recently developed genomic analysis tool SynGO, a database that provides around 3000 expert-curated annotations on 1,112 Synaptic Genes (suppl. table 4) (Koopmans *et al*., 2019). In the target genes for upregulated miRNAs, several were mapped to unique SynGO annotated genes: in lines HD vs pHD, 31 of 363 genes; in lines pHD vs CTR, 25 of 276 genes; and in lines HD vs CTR, 53 of 584 genes (suppl table 5). Similarly, in the target genes for downregulated miRNAs, several were mapped to unique SynGO annotated genes: in lines HD vs CTR, 65 of 751 genes; in lines HD vs pHD, 54 of 408 genes; and in lines pHD vs CTR, 167 of 1440 genes (suppl. table 5).

Since we exposed neuronal cultures to EVs isolated from two specific fibroblast cell lines, one control (fCTR1) and one pHD (fpHD3), we further performed a detailed analysis of the miRNAs identified in these EVs. We found 163 common miRNAs, 89 miRNAs exclusively presented in the EVs from the CTR cell line and 15 miRNAs exclusive to EVs from the pHD cell line (figure 6 A, B). Among the differentially enriched miRNAs, we found 17 upregulated and 7 downregulated in pHD vs CTR (logFC> 2 or logFC< -2, respectively) (figure 6B; supp table 6). To gain further insight into the biological functions of the differentially enriched target genes, we performed GO enrichment analysis for biological process and pathway analysis (suppl. table 7). These genes were mainly involved in biological processes associated with cellular and metabolic pathways.

**Figure 6.**
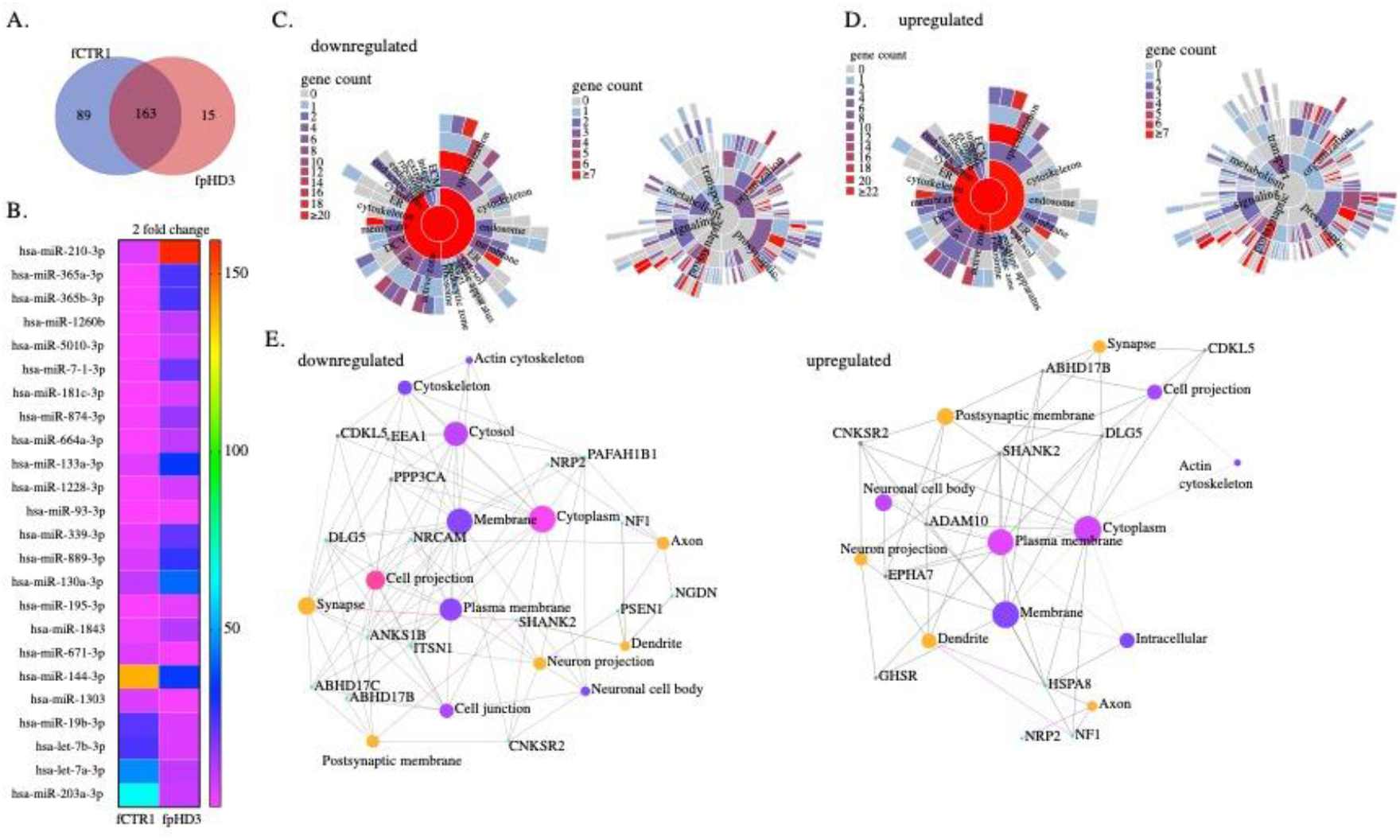
Identification of EVs-miRNAs targeting genes that regulate synaptic function released from HD and CTR fibroblasts. **(A)** Venn diagram showing the number of miRNAs with specific expression for fpHD and fCTR1 cell lines, previously shown to ameliorate abnormal GABAergic function in HD neurons. (B) Heatmap showing the expression of the EVs associated-miRNA from fpHD3 and fCTR1 with a change superior to two-fold. Heatmap colors represent gene expression fold change levels based on the provided color key scale; red for upregulated genes expression levels, pink for downregulated gene expression levels. (C-D) SynGO Cellular Component terms and Biological Process terms are visualized in a sunburst plot for target synaptic genes of upregulated and downregulated EVs-miRNAs. (E) Networks of top enriched cellular components constructed by Network Analyst. Nodes size represents the term enrichment significance by the Bonferroni pos hoc test. The cellular components evaluated are for the set of gene associated with the ontology category GABAergic synapse (GO:0098982).

To better understand if these target genes are involved in synapse biology we applied, as previously, the SynGO analysis to the target genes of downregulated miRNAs. SynGO synaptic ontology can be visualized as sunburst diagrams for synaptic location and function (Figure 6C, D). We found that 333 out of 335 are unique to SynGO annotated genes. Of these, 6 genes for cellular components and 8 for biological processes were significantly enriched at 1% FDR (suppl. table 6) and had functionality in synapse organization, synaptic signaling or chemical synaptic transmission at the presynapse and postsynapse (Figure 6C, suppl. table 6). The SynGO analysis of the target genes for upregulated miRNAs showed that, out of 3014 genes, 310 were mapped to unique SynGO annotated genes and, of these, 6 genes for cellular components and 5 for biological processes were significantly enriched at 1% FDR (Figure 6D, suppl. table 6). Functional GO enrichment analysis indicated that they were mostly involved in synapse organization, synaptic vesicle cycle, synaptic signaling, and trans-synaptic signaling, among others, at the pre and postsynaptic compartment (Figure 6D, suppl. table 6).

To investigate whether the list of target genes of both up- and downregulated miRNAs present in EVs that are involved in synaptic functions could be related with inhibitory synapses (GO:0060077), we compared our results with a list of validated synaptic genes predicted to be expressed in GABAergic synapses (suppl. table 7). We identified several genes potentially targeted by upregulated miRNAs in fpHD that are involved in postsynapse organization (N=22; GSEA p=0.013), regulation of postsynapse organization (N=35; GSEA p=0.05), regulation of translation at synapse, modulating synaptic transmission (N=35; GSEA p=0.08) and regulation of synapse organization (N=7; GSEA p=0,14) (Figure 6E, suppl. table 7).

To date, a few studies have been published reporting brain-specific miRNAs involved in modulating the expression of genes related to GABAergic synaptic function. We first undertook a systematic search for all published studies that identified miRNAs and gene targets that modulate GABAergic synapses and compared them to our miRNAs found in fpHD3- and fCTR1-derived EVs. We identified several dysregulated miRNAs in fpHD3 cells (vs fCTR1) that were described to disrupt GABAergic synapses particularly by modulating GABA receptors or assembly of synapses (Table 2). Some miRNAs were only found in a particular population, CTR or pHD, highly suggesting their involvement in the dysfunction of GABAergic synapses (Table 2). Taken together, these findings show that EVs secreted by cells from HD patients can carry different miRNAs than those carried by EVs from CTR cells, the former being associated with synaptogenesis dysregulation.

**Table 2.**
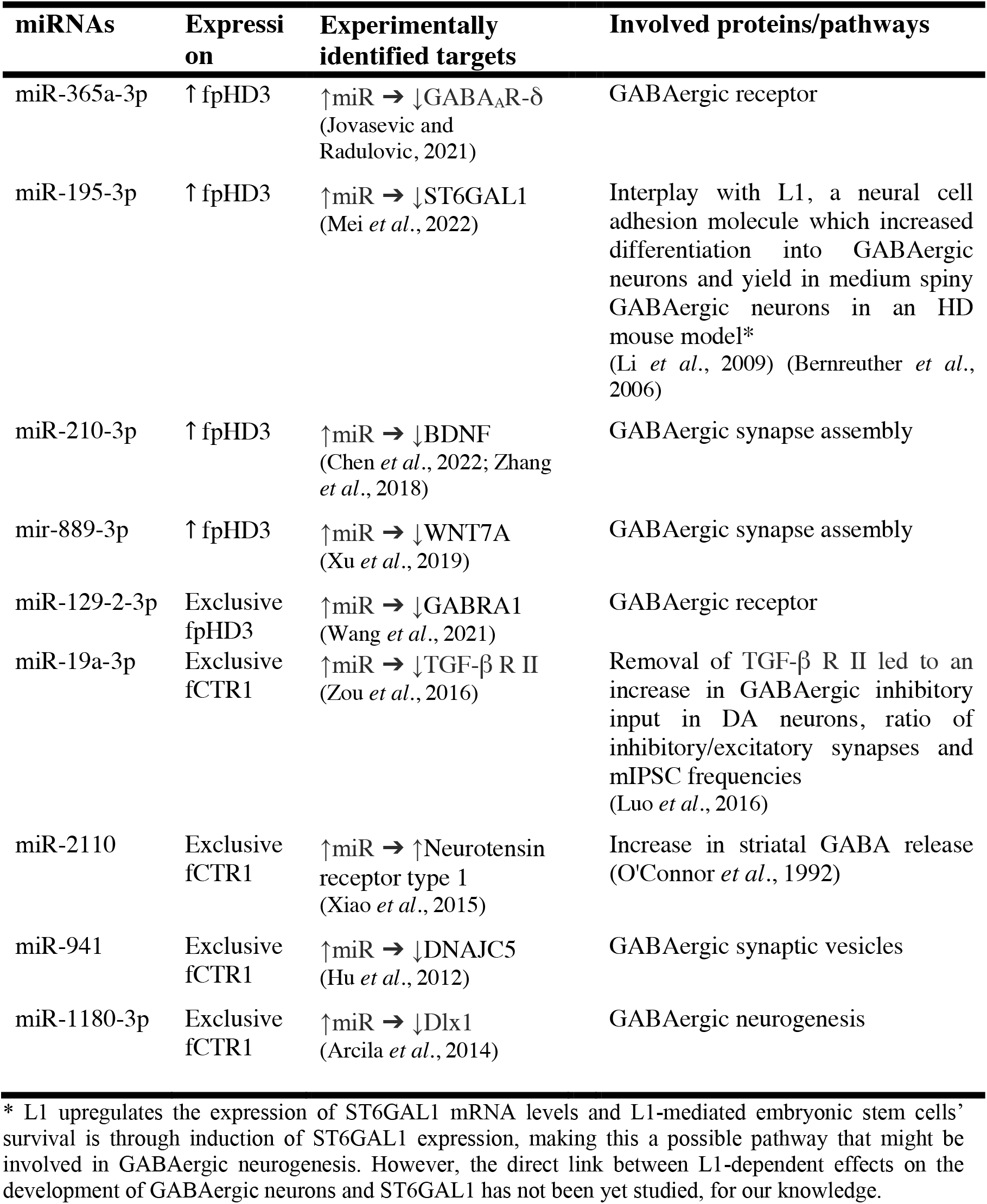
MicroRNAs significantly up- and down-regulated or exclusive of fpHD3- and fCTR1-derived EVs with previously validated targets that modulate GABAergic synapses.

## Discussion

Functional and molecular analyses of GABAergic inhibitory currents in the striatum of HD were associated with degeneration in this structure (Garret *et al*., 2018). Here, we demonstrate that iPSCs reprogrammed from fibroblasts of HD patients can be differentiated into MSN in an equivalent time course as controls, using a previously described protocol (Beatriz *et al*., 2022; Delli Carri *et al*., 2013; Lopes *et al*., 2022). However, the HD-derived cell lines exhibited reduced efficiency in GABAergic striatal neuron specification, as shown by the lower levels of MAP2 and DARPP32 expression, and decreased expression of inhibitory synaptic markers in mature striatal-like neurons, associated to a delayed electrophysiological maturation phenotype. We also observed a lower whole-cell GABAR-mediated current density, mIPSC frequency and density of GABAergic synapses in HD-derived neurons, showing a dysfunction of the GABAergic system compared to CTR-derived neurons. Interestingly, while the treatment of HD-cells with EVs secreted by CTR fibroblasts restored GABAR-mediated current and the inhibitory synaptic density, the incubation of CTR-cells with EVs from HD fibroblasts decreased both GABAR-mediated current density and GABAergic synaptic density. Analysis of the miRNA content in EVs identified predicted target genes with functions at GABAergic synapses that may play a protective role through rescuing of synaptic dysfunction.

Human iPSC systems for modeling HD phenotypes have provided evidence of disturbances in early developmental processes, including altered expression of genes related to neurodevelopmental pathways and synaptic homeostasis (Consortium, 2017; Lopes *et al*., 2016; Lopes et al., 2020). The HD iPSC Consortium showed that more than 50% of differentially expressed genes in HD-derived cell lines were associated with nervous system development and function. Moreover, others identified dysregulated pathways relevant to neuronal development and maturation, included axonal guidance, Wnt signaling, Ca^2+^ signaling (voltage-gated calcium channel subunits, plasma membrane Ca2+ ATPase, CAMKII, CALM and CREB), neuronal CREB signaling, and glutamate and GABA signaling (Consortium, 2017). Several genes with altered expression are related to GABA synthesis, release, reuptake or degradation, suggesting that GABAergic neurons may display increased vulnerability to mHTT cytotoxicity. A delayed electrophysiological maturation phenotype in HD iPSC-derived neurons, associated to altered developmental signatures and reduced number of neural progenitor cells, has been reported in both monolayer neuronal cultures and 3D organoids (Conforti *et al*., 2018; Mattis *et al*., 2015; Mehta *et al*., 2018). Specifically, altered neuronal maturation in cultures of HD iPSC-derived striatal progenitors has been shown, with decreased levels of MAP2 compared to CTRs similarly to what we observed in this work, although there was also a decreased expression of β III-tubulin in HD-cultures vs CTR-cells contrarily to our study (Conforti *et al*., 2018). This discrepancy could maybe be explained by the different time points used to assess β III-tubulin levels. Additionally, we observed that the differentiation into MSN revealed to be lower in HD-cultures by assessing the diminished levels of DARPP32 expression compared to CTRs as previously demonstrated (Conforti *et al*., 2018). Although other authors have showed that HD striatal cultures exhibit more neuronal progenitors than CTR counterparts, they did not find lowered levels of MAP2 nor DARPP32 (Mattis *et al*., 2015). Differentiated HD cortical MAP2+-neurons showed to have decreased neurite length and downregulation of calcium-gated channels, ultimately showcasing alterations in neuronal morphology (Mehta *et al*., 2018). Our data show that HD-derived iPSCs can generate electrophysiologically mature striatal-like neurons that, however, display a more depolarized resting membrane potential and greater excitability, probably due to dysfunctional inwardly rectifying K+ channels, as also reported in other studies using HD mice models (Tong *et al*., 2014; Zhang *et al*., 2018b).

The ionotropic GABA_A_ and glycine receptors, clustered by Gephyrin, were demonstrated to be altered in both HD mouse models and HD patients (Cepeda *et al*., 2007; Dargaei *et al*., 2019; Hsu *et al*., 2018). The underlying molecular mechanisms targeting the GABA_A_ergic system in the context of HD are still unknown, but evidence suggests that mHTT alters the transcription of genes and protein expression (e.g. GABA_A_R and KCC2) through interactions with transcriptional activators and repressors, but also other factors such as excitotoxicity due to impaired homeostasis of extracellular glutamate, neuroinflammation caused by the interaction of mHTT with astrocytes and microglia and reduced GABA synthesis by astrocytes (Dargaei *et al*., 2019; Pribiag and Stellwagen, 2013; Skotte *et al*., 2018; Stellwagen *et al*., 2005; Yuen *et al*., 2012). Additionally, evidence has showed that GABA_A_R subunits (α 1, α 2) are specifically altered in striatal MSN of HD mouse models R6/1 and HdhQ111 (Du *et al*., 2017). In this study, we used peripheric EVs derived from human fibroblasts to treat differentiated neuronal cultures and observed the effects on GABAergic function and synaptic density. Previously, it has been demonstrated that human EVs are able to cross the blood-brain barrier, acting as messengers between the peripheric blood system and central nervous system (Matsumoto *et al*., 2017). A study using Parkinson disease patients’ blood showed that EVs derived from erythrocytes can have an impact on microglia by transporting alpha-synuclein across the blood-brain barrier (Matsumoto *et al*., 2017).

Here, we demonstrate that several miRNAs involved in modulating key target genes related to inhibitory synapse function are upregulated in EVs isolated from HD fibroblasts, suggesting that the functional/developmental impairment identified in neurons treated with these EVs could be, at least partially, mediated by their miRNA cargo. Several hypothetical downregulated genes targeted by the upregulated miRNAs identified in HD-EVs are known to regulate GABAergic synapse-related proteins (Table 2). In agreement, the metalloproteinase ADAM10 shedding enzyme, that is a known target for upregulated miRNAs found in HD-EVs, is a central protein for the development of the nervous system (Jorissen *et al*., 2010). Indeed, the loss of ADAM10 is linked to defects in neuronal connectivity and the postnatal neuron-specific disruption of ADAM10 in the brain leads to impaired learning, defects in long-term potentiation and seizures (Kuhn *et al*., 2016). Another identified target was the NF1 gene, whose the protein expression levels are significantly higher in inhibitory neurons than in excitatory neurons. The downregulation of NF1 expression influences both GABA concentration and GABAA receptor density, suggesting a pre- and postsynaptic involvement (Violante *et al*., 2016). Another protein that is highly expressed in GABAergic neurons is MeCP2. In MeCP2 KO mice, the GABAergic synaptic transmission is strongly depressed (Medrihan *et al*., 2008). Mutations in the MeCP2 gene that cause Rett syndrome, a prevalent neurodevelopmental disorder, result in an excitatory–inhibitory imbalance (Zhang *et al*., 2010). Our data demonstrate that several gene candidates are predicted to affect multiple processes and contribute for the reduced GABAergic synaptogenesis. Consistent with this, miRNAs identified in our study as being upregulated in HD-EVs were previously validated as targeting proteins/pathways regulating inhibitory synapses (Jovasevic and Radulovic, 2021; Wang *et al*., 2021; Zhang *et al*., 2018a).

Our data from CTRs-EVs supports the upregulation of some particular miRNAs with previously validated targets that modulate GABAergic synapses (eg. miR-19a-3p, miR-211, miR-1180-3p, miR-941) (Arcila *et al*., 2014; Bernreuther *et al*., 2006; Chen *et al*., 2022; Hu *et al*., 2012; Jovasevic and Radulovic, 2021; Li *et al*., 2009; Luo *et al*., 2016; Mei *et al*., 2022; O’Connor *et al*., 1992; Wang *et al*., 2021; Xiao *et al*., 2015; Xu *et al*., 2019; 2018a; Zou *et al*., 2016) (Table 2). Interestingly, the cysteine string protein, encoded by the *DNAJC5* gene, resides in presynaptic terminals, clathrin-coated vesicles and neuroendocrine secretory granules and is involved in neurotransmitter release and most importantly has been linked to Huntington’s and Parkinson’s diseases (Chandra *et al*., 2005; Miller *et al*., 2003; Ruiz *et al*., 2008). Additionally, EVs are also implicated in the development of neurodegenerative diseases, particularly proteinopathies, as they carry and spread disease-associated proteins (Beatriz *et al*., 2021). In accordance, our data indicate HD-EVs carry mHTT, as previously reported (Beatriz *et al*., 2022; Hong *et al*., 2017). Our previous data have demonstrated that mitochondria-lysosomal axis dysregulation in HD enhances EVs release along with mitochondrial DNA and proteins cargo (Beatriz *et al*., 2022).

In a recent study, Sharma et al. validated the role of EVs in synaptogenesis and neural circuit development and hypothesized that the effect could be mediated by the proteomic cargo of these vesicles (Sharma *et al*., 2019). Authors reported increased neuronal proliferation and synaptic density in human-derived differentiated neuronal cultures with MECP2 loss-of-function upon incubation with EVs from isogenic CTRs (Sharma *et al*., 2019). Moreover, addition of CTR-EVs to human-derived neurospheres with MECP2 loss-of-function resulted in an increase in neural activity (Sharma *et al*., 2019). Furthermore, injection of CTR NSC-derived EVs in Alzheimer disease (AD) mice led to an increase in GAP43, synaptophysin and PSD-45 synaptic markers in the cortex when compared to vehicle-injected animals (Li *et al*., 2020). In epileptic mice, the use of EVs isolated from human bone marrow-derived mesenchymal stem cells improved glutamatergic and GABAergic loss in the hippocampus (Long *et al*., 2017), and mice that suffered transient ischemia showed ameliorated hippocampal long-term potentiation and synaptic transmission by recovering field excitatory postsynaptic potential (Deng *et al*., 2017). Moreover, the analysis of the content of stem cells-EVs (human iPSC-derived NSC and adipose-derived mesenchymal stem cells) demonstrated that these vesicles are carriers for proteins and miRNAs associated to neuroprotection and synaptogenesis as is the case of agrin and neuroplastin (Ma *et al*., 2020; Upadhya *et al*., 2020). Additionally, the injection of such EVs successfully led to increased hippocampal neurogenesis in WT rat and AD mouse brains, respectively (Ma *et al*., 2020; Upadhya *et al*., 2020). Our study shows that administration of CTR-EVs might provide a therapeutic strategy to ameliorate neuronal function and synaptic maintenance, specifically in early stages of HD, before a major degeneration of GABAergic neurons. Understanding EV-mediated molecular mechanisms of signaling underlying this functional impact is needed to allow the exploitation of their therapeutic potential.

In conclusion, our data demonstrate for the first time that EVs from HD patients carry cell-derived bioactive molecules that have an impact on cellular functions, particularly in neurodegenerative disease models with synaptopathy as a central feature. Additionally, the EVs-mediated reversal of the pathological phenotype provides a rational basis for further study investigating their use in therapeutic applications.

## Acknowledgments

We acknowledge Dr. Paulo Oliveira for critical review of the manuscript, Dr. Mónica Zuzarte for TEM analysis at the ‘Laboratório de Bio-imagem de Alta Resolução’, Faculty of Medicine of the University of Coimbra, Dr. Teresa Rodrigues and Dr. Henrique Girão from the Institute of Biomedical Imaging and Life Sciences (IBILI), Faculty of Medicine, University of Coimbra for NTA equipment.

## CRediT authorship contribution statement

Carla Lopes, A. Cristina Rego designed this study. Margarida Beatriz, Ricardo Rodrigues, Rita Vilaça, Carla Lopes: performed the experiments. Conceição Egas: performed RNA sequencing and analysis. Margarida Beatriz, Ricardo Rodrigues, Rita Vilaça, Carla Lopes: analysed the data, Carla Lopes, Margarida Beatriz: drafted the initial manuscript. Margarida Beatriz, Ricardo Rodrigues, Rita Vilaça, Conceição Egas, Paulo Pinheiro, George Q. Daley, Thorsten M. Schlaeger, A. Cristina Rego, Carla Lopes: revised this manuscript. all authors read and approved the final manuscript. Carla Lopes: secured funding.

