## Supplementary figures and images for "Extracellular vesicles improve GABAergic transmission in Huntington’s disease iPSC-derived neurons"

### Supplementary figure 1

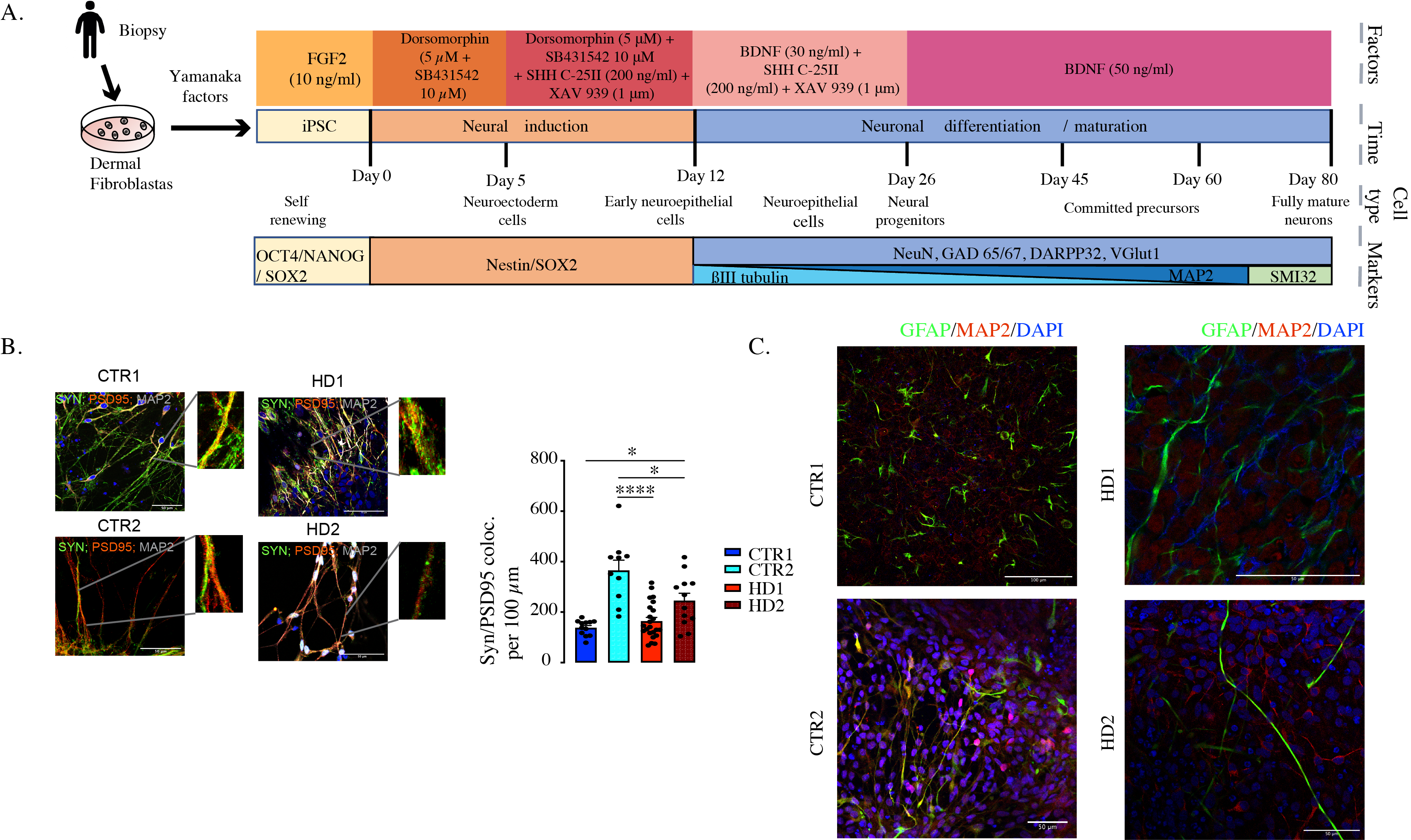

### Supplementary figure 2

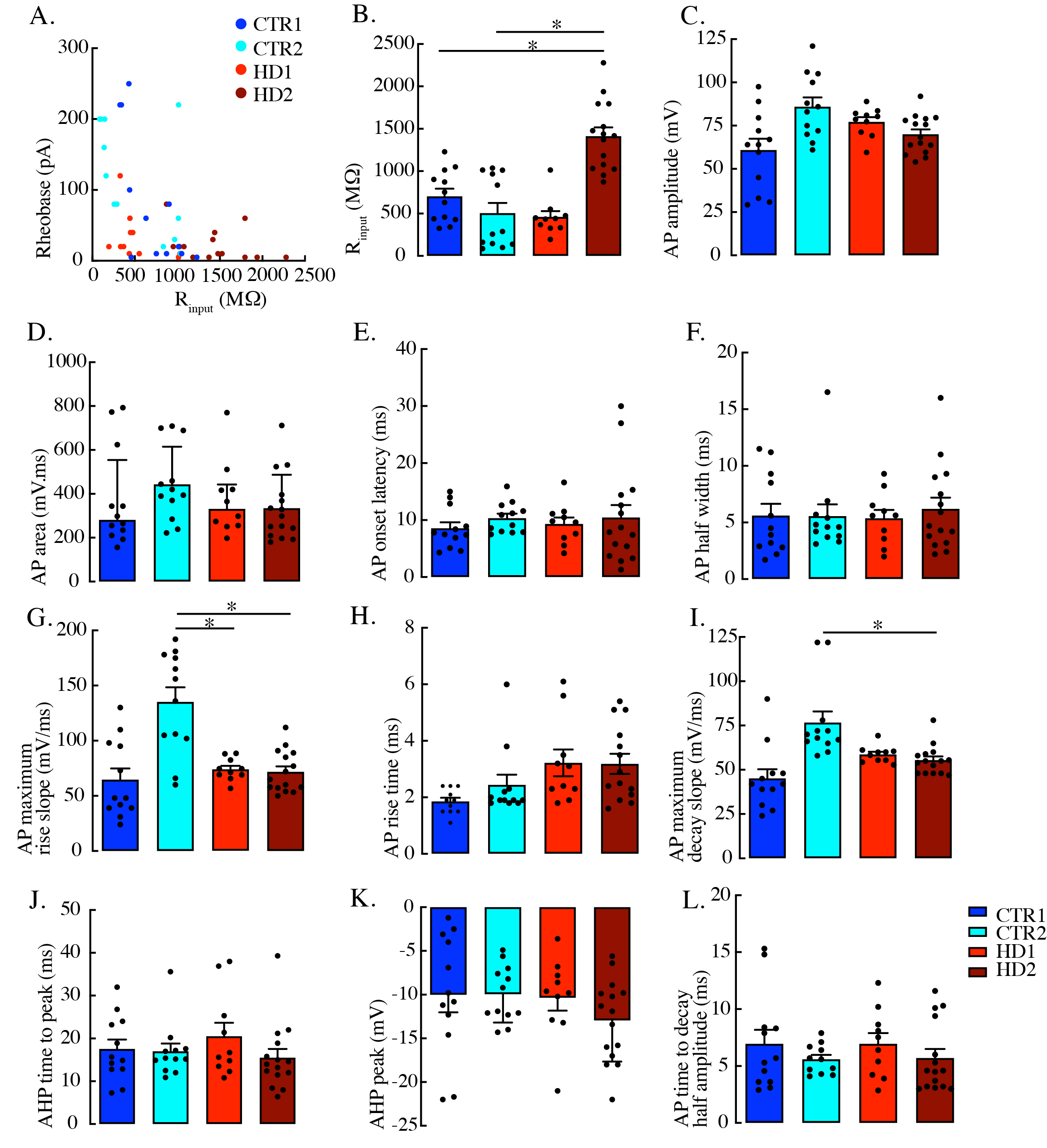

### Supplementary figure 3

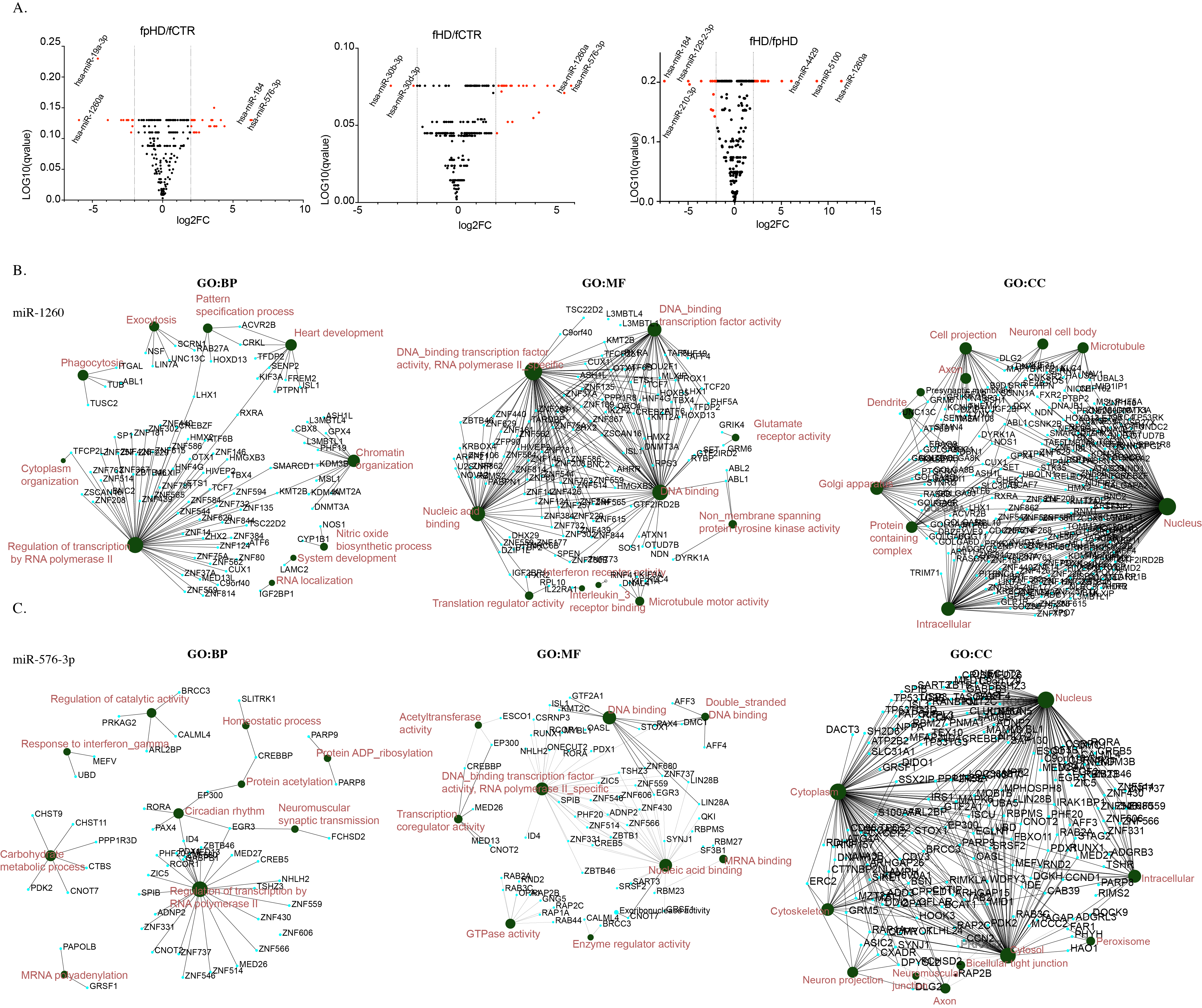
