## Supplementary table 1 for "Extracellular vesicles improve GABAergic transmission in Huntington’s disease iPSC-derived neurons"

**Supplementary Table 1.** Metrics per sample along the different analysis steps.

| **Metrics /**  **samples** | **Total Reads**  **sequenced (#)** | **Total Reads after**  **QC (#)** | **Total Reads after**  **rRNA and tRNA**  **removal (#)** | **Reads mapped**  **to miRBase (#)** |
| --- | --- | --- | --- | --- |
| fCTR1 | 13,347,636 | 9,963,425 | 4,342,630 | 1,103,931 |
| fCTR2 | 13,152,667 | 5,324,116 | 2,492,835 | 687,854 |
| fCTR3 | 11,675,829 | 7,113,831 | 4,452,039 | 2,081,559 |
| fpHD1 | 13,392,005 | 8,171,010 | 5,260,904 | 404,498 |
| fpHD2 | 13,815,452 | 7,939,090 | 5,747,350 | 4,890,054 |
| fpHD3 | 11,245,268 | 7,058,606 | 4,499,043 | 238,631 |
| fHD1 | 11,121,267 | 4,307,082 | 1,821,026 | 1,077,743 |
| fHD2 | 9,725,087 | 4,574,519 | 3,095,192 | 2,576,082 |
| fHD3 | 10,327,642 | 6,174,631 | 3,632,983 | 2,856,454 |

Total Read Sequences – number of raw sequences obtained from the NextSeq sequencing platform; Total Reads after QC – number of reads after removing adapters and 4Ns introduced in library preparation; Total Reads after rRNA and

tRNA removal against RFam – number of reads not corresponding to rRNA and tRNA; Reads mapped to MIRBase – number of reads mapped to MIRBase, version 22.
