## Supplementary data for "Extracellular vesicles improve GABAergic transmission in Huntington’s disease iPSC-derived neurons"

**Figure S1.** Differentiation protocol for striatal-like neurons.

(A) The iPSC markers include OCT4 and NANOG, and SOX2 is still present as neuroectoderm cells start to appear during neuronal induction together with Nestin. Neuronal differentiation gives origin to neuroepithelial cells and βIII-tubulin is present in immature neurofilaments whereas MAP2 is present as long as neurofilaments mature. Cultures of neuronal progenitors with a striatal lineage include NeuN, GAD 65/67, DARPP32 and VGlut1 as markers. SMI32 is present in fully mature neurons at day 80 of differentiation. (B) Mature striatal-like neurons (MAP2^+^) presented presynaptic (synaptophysin) and excitatory postsynaptic (PSD95) markers; colocalization of synaptophysin with PS95 in mature striatal-like neurons. (C) Microglia marker (GFAP) is present in neuronal (MAP2+) cultures at day 80. Bar plots represent mean±S.E.M. Statistical analysis: one-way ANOVA followed by Bonferroni multiple comparisons test: * p<0.05, **** p< 0.0001.

**Figure S2.** Electrophysiological properties of HD-derived neurons and CTRs.

Bar plots showing (A) Rheobase (pA), (B) Input Resistance (MΩ), (C) action potential AP amplitude, (D) AP area onset, (E) AP latency, (F) AP ½-width, (G) AP maximum rise slope, (H) AP rise time, (I) AP maximum decay slope, (J) AP time to decay half amplitude, (K) after-hyperpolarization potential (AHP) time to peak and (L) AHP peak. One-way ANOVA followed by Tukey's multiple comparisons test: * p<0.05.

**Figure S3.** Differentially expressed miRNAs.

(A) Volcano plot of DE miRNAs in EVs isolated from fCTR, fpHD and fHD cell lines as assessed by a microarray analysis. The horizontal axis represents the log2 ratio and the vertical axis represents −log10 (q value). The red color indicates that the expression of the miRNA significantly increased/decreased more than two-fold in EVs from the represented groups. (B-C) Enriched biological processes of miR-1260 family and miR-576-3p gene targets based on overrepresentation analysis (ORA). Gene set nodes are colored based on their enrichment p-value.

**Supplementary Table 1:** Metrics per sample along the different analysis steps

**Supplementary Table 2:** MiRNA differently expressed

**Supplementary Table 3:** miRDB target prediction data (related to figure 5)

**Supplementary Table 4:** Gene Ontology biological process enrichment analysis of differentially expressed genes (up/downregulated) for candidate genes identified through miRDB analysis

**Supplementary Table 5:** Intersection of the gene list from supplementary table 2 with SynGO annotated genes that uses list of all brain expressed genes as a default background set, defined as 'expressed in any GTEx v7 brain tissue'.

**Supplementary Table 6:** miRDB gene target prediction data for fCTR2 and fpHD3 and intersection of the gene list with SynGO annotated genes (related to figure 6)

**Supplementary Table 7:** Intersection of the gene list with SynGO annotated genes for fCTR2 and fpHD3 using the Gene Ontology Consortium database for two categories: GABAergic synapse (GO:0098982) and Synapse assembly (GO:0007416).
